# Endothelial Brg1 fine-tunes Notch signaling during zebrafish heart regeneration

**DOI:** 10.1101/2022.05.18.492445

**Authors:** Chenglu Xiao, Junjie Hou, Fang Wang, Yabing Song, Jiyuan Zheng, Lingfei Luo, Jianbin Wang, Wanqiu Ding, Xiaojun Zhu, Jing-Wei Xiong

**Author notes:** Corresponding authors: Dr. Jing-Wei Xiong, Dr. Xiaojun Zhu, and Dr. Wanqiu Ding. Contributed equally to this work (Chenglu Xiao, Junjie Hou).

## Abstract

Myocardial Brg1 is essential for heart regeneration in zebrafish, but it remains unknown whether and how endothelial Brg1 plays a role in heart regeneration. Here, we found that both *brg1* mRNA and protein were induced in cardiac endothelial cells after ventricular resection, and endothelium-specific over-expression of dominant-negative *Xenopus* Brg1 (DN-xBrg1) inhibited myocardial proliferation and heart regeneration and increased cardiac fibrosis. RNA-seq and ChIP-seq analysis revealed that the endothelium-specific over-expression of DN-xBrg1 changed the levels of H3K4me3 modifications in the promoter regions of the zebrafish genome and induced abnormal activation of Notch family genes upon injury. Mechanistically, Brg1 interacted with lysine demethylase 7aa (Kdm7aa) to fine-tune the level of H3K4me3 within the promoter regions of Notch family genes and thus regulated Notch gene transcription. Together, this work demonstrates that the Brg1-Kdm7aa-Notch axis in cardiac endothelial cells, including the endocardium, regulates myocardial proliferation and regeneration *via* modulating the H3K4me3 of the Notch promoters in zebrafish.

## Introduction

The high mortality and morbidity of myocardial infarction is of public concerns worldwide. The loss of cardiomyocytes following myocardial infarction and the inadequate self-repair capability of the mammalian heart make it difficult to treat cardiac diseases (Hesse, Welz, & Fleischmann, 2018). As one of the least regenerative organs in the human body, the heart replaces the infarcted myocardium with non-contractile scar instead of new muscles, which is initially beneficial but eventually leads to loss of contraction and function. Although various cell-based and cell-free strategies have been explored to restore infarcted heart function, the efficacy and side-effects such as arrhythmia and immune rejection currently prevent translation to the clinic. The neonatal mouse can regenerate its heart but this ability is lost after 7 postnatal days (Porrello et al., 2011; Sadek & Olson, 2020; Tzahor & Poss, 2017). A number of elegant studies have provided evidence for the underlying mechanisms, but how to efficiently stimulate mammalian heart regeneration remains largely unknown. Unlike mammals, some lower vertebrates such as zebrafish can fully regenerate the heart after injury throughout life (Gemberling, Bailey, Hyde, & Poss, 2013). Dissecting the cellular and molecular mechanisms of zebrafish heart regeneration may provide clues for promoting heart regeneration in mammals.

It is conceivable that cardiomyocyte dedifferentiation and proliferation contribute to heart regeneration in zebrafish (Jopling et al., 2010; Kikuchi et al., 2010). Over the past decades, a number of signaling pathways and transcription factors have been reported to regulate myocardial proliferation and regeneration in zebrafish, including fibroblast growth factor, sonic hedgehog, retinoic acid, insulin-like growth factor, Notch, GATA4, Hand2, NF-kB, and Stat3 (Kikuchi et al., 2011; Pronobis & Poss, 2020; Raya et al., 2003; Zhao, Ben-Yair, Burns, & Burns, 2019; Zhao et al., 2014; Zheng et al., 2021). Retinaldehyde dehydrogenase 2, which produces retinoic acid, is activated in the epicardium and endocardium within hours after injury, and transgenic inhibition of retinoic acid receptors impairs myocardial proliferation (Kikuchi et al., 2011). Conditional inhibition of Notch signaling *via* overexpression of dominant-negative Notch transcriptional co-activator Master-mind like-1 (MAML) in endothelial cells (including the endocardium) decreases myocardial proliferation (Gao, Fan, Zhao, & Su, 2021; Zhao et al., 2019). These studies suggest an essential role of endocardial signaling in regulating myocardial proliferation, but it remains to be addressed how endocardial Notch components are regulated or how endocardial signals regulate myocardial proliferation and regeneration upon injury.

Epigenetic regulation plays an important role in gene expression in various cellular process such as differentiation, proliferation, fate determination, as well as organ regeneration (Duncan & Sanchez Alvarado, 2019; Li & Reinberg, 2011; Martinez-Redondo & Izpisua Belmonte, 2020; Zhu, Xiao, & Xiong, 2018). Epigenetic regulation is in general defined as controlling gene expression beyond the DNA sequence itself, consisting of histone modifications, DNA/RNA modifications, non-coding RNAs, and chromatin remodeling complexes (Oyama, El-Nachef, Zhang, Sdek, & MacLellan, 2014). The SWI/SNF (SWItch/Sucrose Non-Fermentable)-like complex, a member of the ATP-dependent chromatin-remodeling complex family, uses energy from ATP hydrolysis, regulates gene transcription by rearranging nucleosome positions and histone-DNA interactions, and thus facilitates the transcriptional activation or repression of targeted genes (Ho & Crabtree, 2010). We previously reported that its central subunit, brahma-related gene 1 (Brg1 or Smarca4), had critical function in zebrafish heart regeneration by interacting with DNA (cytosine-5-)-methyltransferase 3 alpha b to modify DNA methylation of the cyclin-dependent kinase inhibitor 1C promoter (Xiao et al., 2016). We found that *brg1* was not only induced in cardiomyocytes but also in cardiac endothelial cells, including the endocardium, during myocardial regeneration (Xiao et al., 2016). In this work, we investigated how endothelial Brg1 played a role in zebrafish heart regeneration. Inhibition of Brg1 *via* dominant-negative (DN)-xBrg1 in cardiac endothelial/endocardial cells decreased myocardial proliferation and heart regeneration, and Brg1 interacted with the histone demethylase Kdm7aa (lysine (K)-specific demethylase 7Aa) to regulate Notch receptor gene expression upon injury. Together, this work presents the first evidence, to our knowledge, that the Brg1-Kdm7aa axis fine-tunes Notch signaling in cardiac endothelium and endocardium during heart regeneration.

## Results

### Endothelial Brg1 is required for heart regeneration in zebrafish

Our previous work has shown that both global and cardiac-specific inhibition of Brg1 results in impaired myocardial proliferation and regeneration, while global inhibition of Brg1 leads to more severe cardiac fibrosis than its myocardium-specific inhibition (Xiao et al., 2016). In addition to elevated expression in the injured myocardium, Brg1 was also induced in other cardiac cells including endothelial cells during heart regeneration. To evaluate Brg1 expression in endothelial cells during zebrafish heart regeneration, we used immunofluorescence staining (Fig. 1A, B) and RNAscope *in situ* hybridization (Fig. 1C, D) to determine whether Brg1 was induced in endothelial cells upon ventricular amputation. Consistent with our previous report, Brg1 protein was co-localized with Tg(*fli1*:nucEGFP)-positive endothelial cells in the injury site at 7 days post-amputation (dpa) (Fig. 1A, B). Moreover, RNAscope staining revealed that *brg1* mRNA was elevated and partially overlapped with *kdrl*-positive endothelium at 3 dpa (Fig. 1C, D). We then turned to tamoxifen-induced endothelium-specific inhibition of Brg1 with the transgenic strains Tg(*ubi*:loxp-DsRed-STOP-loxp-DN-xBrg1; *kdrl*:CreER) (Xiao et al., 2016; Zhan et al., 2018) to address whether Brg1 had a function in endothelial cells during regeneration. We found that endothelium-specific over-expression of DN-xBrg1 resulted in abnormal cardiac fibrosis (Fig. 1E, F) and compromised myocardial regeneration (Fig. 1G, H) at 30 dpa as well as decreased proliferating cardiomyocytes at 7 dpa (Fig. 1I-K).

**Figure 1.**
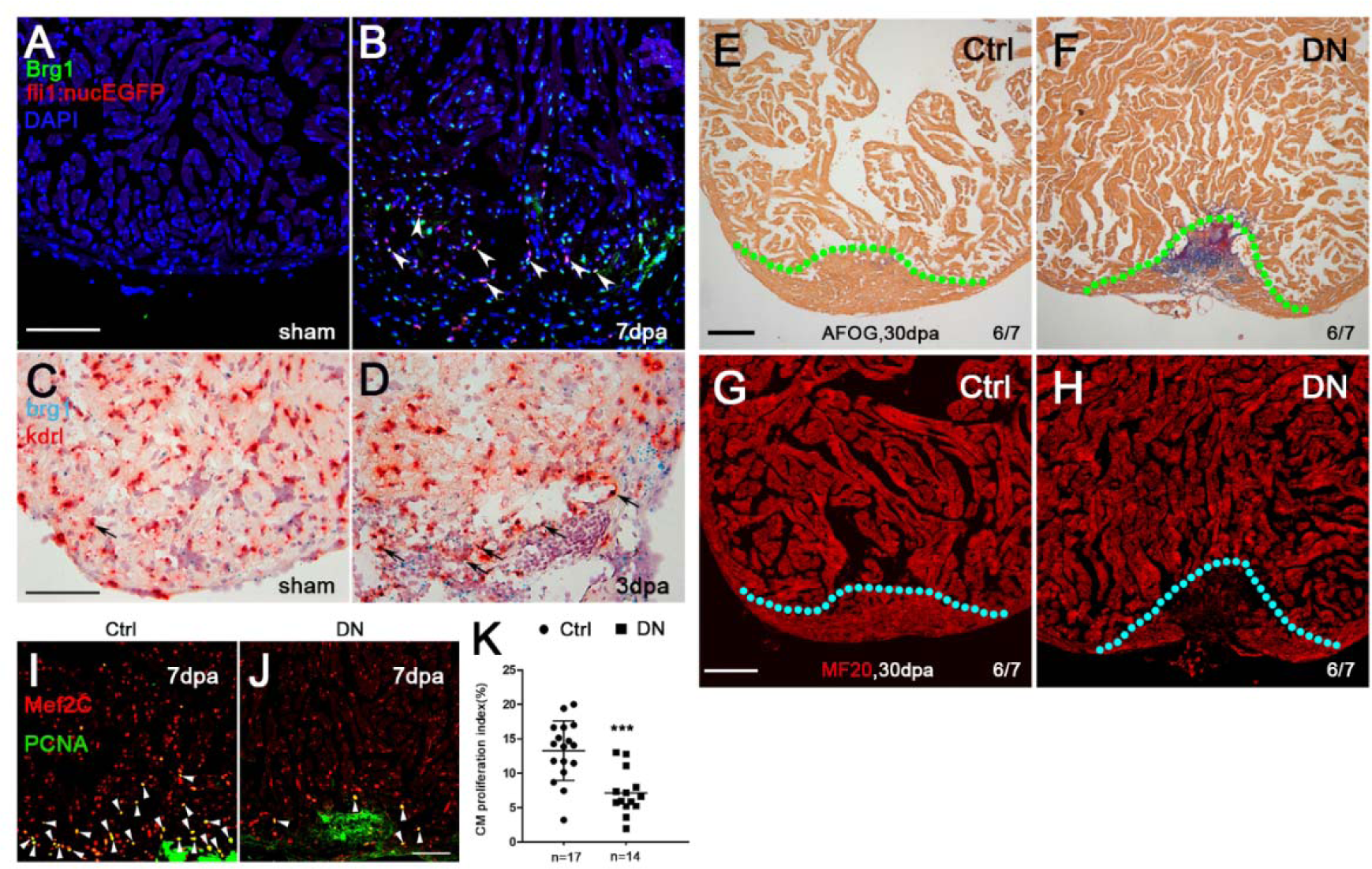
Inhibition of endothelial Brg1 impairs myocardial proliferation and regeneration. **(A, B)** Immunofluorescence staining of Brg1 and EGFP on paraffin sections of Tg(*fli1*:nucEGFP) transgenic hearts from sham-operated (A) and injured zebrafish hearts (B) at 7 dpa (arrowheads, Brg1- and EGFP-positive endothelial cell nuclei). **(C**, **D)** RNAscope *in situ* hybridization of *brg1* and *kdrl* probes in frozen sections from sham-operated (C) and injured hearts (D) at 3 dpa (arrows, *brg1*- and *kdrl*-positive endothelial cells). **(E**–**H)** Representative images of Acid Fuschin-Orange G (AFOG) staining (E, F) and immunofluorescence with anti-myosin heavy chain (MF20) (G, H) of heart sections from control siblings Tg(*ubi*:loxp-DsRed-STOP-loxp-DN-xBrg1) (Ctrl) and endothelium-specific dominant-negative brg1 mutants Tg(*ubi*:loxp-DsRed-STOP-loxp-DN-xBrg1; *kdrl*:CreER) (DN) at 30 dpa, noting that, compared with robust regenerated myocardium and rare cardiac fibrosis in Ctrl group (E, G), the DN group failed to regenerate the myocardium (H) and had evident fibrin (red) and collagen (blue) deposition (F). Dashed lines mark the resection traces. N numbers indicate biological replicates. **(I**, **J)** Immunostaining of representative heart sections at 7 dpa identified cardiomyocyte nuclei (Mef2C^+^) and nuclei undergoing DNA replication (PCNA^+^). Noting fewer proliferative cardiomyocytes (Mef2C^+^/PCNA^+^) in the DN group than in the Ctrl group. Arrowheads, Mef2C^+^/PCNA^+^ proliferating cardiomyocytes. **(K)** Statistical analysis of experiments as in I and J (CM, cardiomyocyte; n = 17 biological replication for Ctrl group and 14 biological replications for DN group; data are the mean percentage ± s.e.m.; ***p <0.001, unpaired *t*-test). Scale bars, 100 μm. **Figure 1-source data 1.** Source images for Figure 1E-J. **Figure 1-source data 2.** Source data for Figure 1K.

Using RNAscope *in situ* hybridization, we also found that endothelium-specific inhibition of Brg1 interfered with the formation of *kdrl*-positive endothelial cells (Figure 1-figure supplement 1A-C) and *coronin1a*-positive leukocytes (Figure 1-figure supplement 1D-F) while having no effect on *tcf21*-positive epicardium (Figure 1-figure supplement 1G-I) in DN hearts compared with Ctrl sibling hearts at 7 dpa in the presence of 4-hydroxytamoxifen (4-HT). Taken together, these data demonstrate that endothelial Brg1 is required for myocardial proliferation, angiogenesis, and leukocyte recruitment but not for epicardium formation during heart regeneration.

### Endothelium-specific inhibition of Brg1 changes the levels of H3K4me3 in the promoter regions of zebrafish genome

To decipher the molecular action of endothelial Brg1, we used RNA-seq analysis to search for Brg1-regulated genes during heart regeneration. We applied Tg(*kdrl*:eGFP) to label cardiac endothelial cells including the endocardium, and achieved endothelium-specific over-expression of DN-xBrg1 by using the compound zebrafish line consisting of Tg(*ubi*:loxp-DsRed-STOP-loxp-DN-xBrg1; *kdrl*:CreER; *kdrl*:eGFP) (defined as DNK), while we used Tg(*ubi*:loxp-DsRed-STOP-loxp-DN-xBrg1; *kdrl*-eGFP) as control (CtrlK) in the presence of 4-HT starting at 3 days before ventricular resection. The *kdrl*:eGFP endothelial cells, which were sorted by fluorescence-activated cell sorting (FACS) from CtrlK and DNK hearts at 7 dpa, were subjected to RNA-seq analysis, and differentially-expressed genes were identified (Fig. 2A). Compared with CtrlK group, we found 1,163 up-regulated genes and 1,266 down-regulated genes in DNK group (Fig. 2A; Figure 2-source data 1). Further bioinformatics analyses of these genes revealed that receptor activity related genes were among the top-affected leads, in which the Notch signaling component *notch2* was strongly induced in DNK group (Fig. 2A; Figure 2-figure supplement 1A). Other genes related to mitosis and cell-cycle were down-regulated, while the genes related to collagen and fibronectin were up-regulated in DNK group compared to CtrlK sibling group (Figure 2-figure supplement 1B).

**Figure 2.**
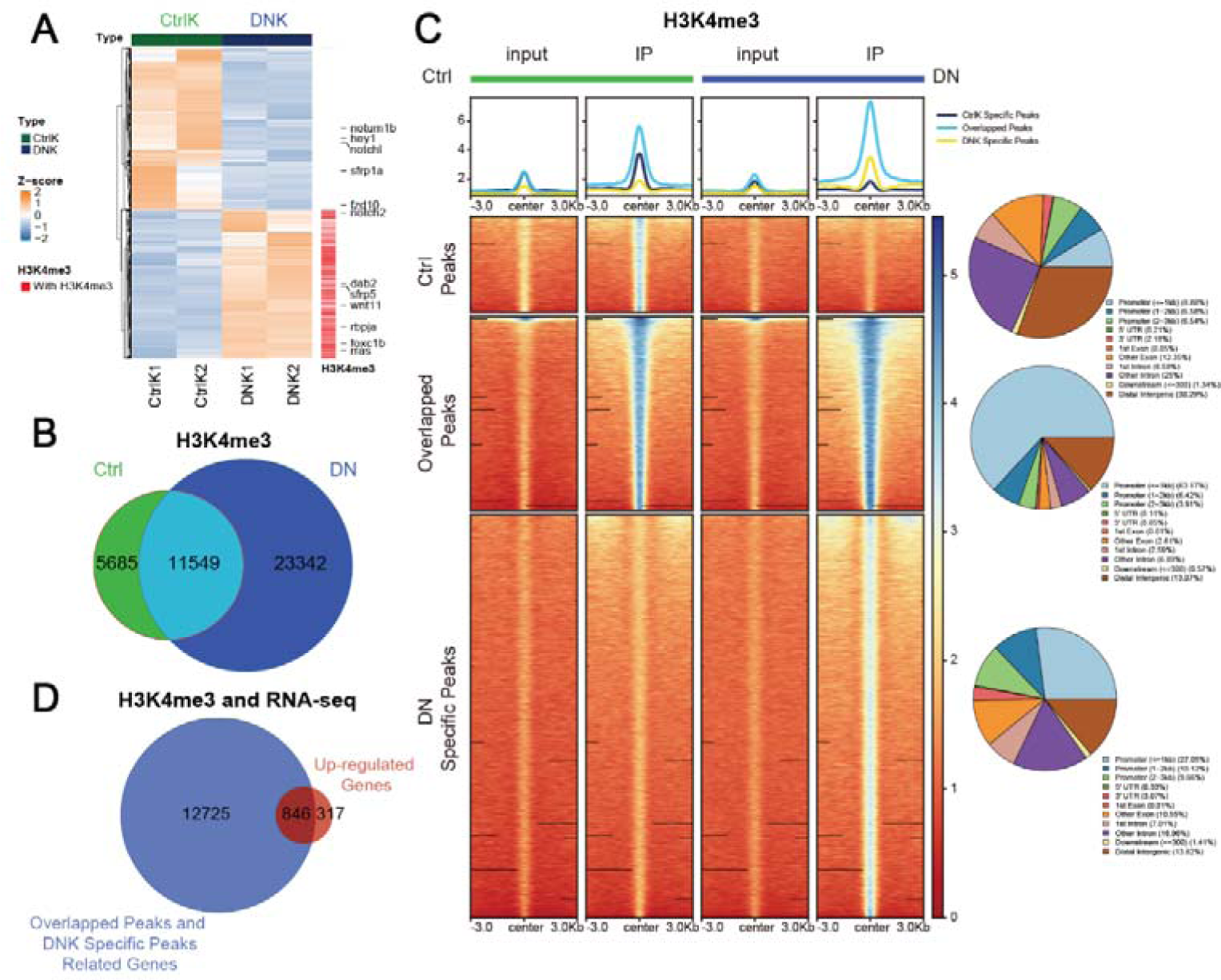
Endothelium-specific inhibition of Brg1 changes the levels of H3K4me3 in the promoter regions of the zebrafish genome. **(A)** Heat map displaying Z-score normalized gene expression for differentially-expressed genes between *kdrl*-eGFP positive endothelial cells from dominant-negative Brg1 groups (DNK1 and DNK2) and control groups (CtrlK1 and CtrlK2). FPKM value (The Fragments Per Kilobase of transcript per Million mapped reads) of each gene was normalized using Z-scores across samples. Columns represent individual samples (two biological replicates for each group); rows represent differentially-expressed genes ordered by hierarchical clustering. Labeled genes are part of the differentially-expressed Notch signaling genes. The up-regulated genes in DNK group that are labelled with ‘red color’ had H3K4me3 peaks in their promoters. **(B)** Venn plot representing the intersection of H3K4me3 peaks between Ctrl and DN groups. **(C)** Heatmaps and summary plots displaying the signal profile of normalized read coverage around three categories of H3K4me3 peaks across different samples (inputs and IP samples in Ctrl and DN groups, respectively). The read coverage was normalized to 1x sequencing depth in all samples. Each row of heatmap represents one peak, with coverage plotted across the 3kb surrounding the peak summit. H3K4me3 peaks are classified into three categories: Ctrl Specific Peaks represent peaks specifically in Ctrl group; Overlapped Peaks represent peaks overlapped between Ctrl and DN groups; DN Specific Peaks represent peaks specifically in DN group. The genomic distribution for three types of peaks is presented with pie charts on the right side. **(D)** Venn plot representing the intersection between genes with promoters marked by Overlapped Peaks and DN Specific Peaks and genes that are differentially upregulated in DNK group. “*notch2*” is indicated in the overlapped gene list. **Figure 2-source data 1**. FPKM values for all differential expressed genes in each condition shown in Figure 2A. **Figure 2-source data 2**. Raw H3K4me3 peak files called by MACS2 in two conditions for Figure 2B and peak files for three categories peaks shown in Figure 2C.

It is well recognized that Brg1 is involved in both gene activation and repression through interacting with epigenetic modifiers and influencing histone modifications at the targeted gene promoters (Menon, Shibata, Mu, & Magnuson, 2019). And previous studies have established that the nucleosomes with histone H3 Lysine 4 trimethylation (H3K4me3) are mainly associated with the promoter regions of active transcription (Vastenhouw et al., 2010; W. Zhu, Xu, Wang, & Liu, 2019). Therefore, we examine whether endothelial-specific overexpression of DN-xBrg1 has effect on the level of the histone marker H3K4me3 in the zebrafish genome. Genome-wide ChIP-seq analyses of Ctrl and DN amputated ventricles at 7dpa using H3K4me3 antibody revealed that, in addition to 11,549 overlapping H3K4me3 peaks between the Ctrl and DN groups, more H3K4me3 peaks emerged in DN group, suggesting that inhibition of Brg1 enhanced H3K4me3 modifications (Fig. 2B). Peaks were then divided into three categories according to the Venn plot, namely Ctrl Specific Peaks in Ctrl group, Overlapped Peaks representing peaks overlapped between Ctrl and DN groups, and DN Specific Peaks representing peaks specifically in DN group. Heatmaps and summary plots of H3K4me3 ChIP-seq signals in 3 kb surrounding the peak summits displayed slightly stronger Ctrl Specific Peaks signals in Ctrl group, while increased Overlapped Peaks signals and DNK Specific Peaks signals in DN group (Fig. 2C).

Moreover, genomic distribution analysis for three categories of peaks revealed that peaks with increased signals in DN group were more concentrated in the promoter region than that with decreased peak signals (Fig. 2C), suggesting that endothelial Brg1 inhibition led to elevated levels of H3K4me3 in the promoter regions. We then examined the correlation of differentially expressed genes from RNA-seq and H3K4me3 modification levels. We analyzed the overlapping genes by comparing up-regulated genes in DNK group with the genes that their promoters were marked by Overlapped Peaks and DN Specific Peaks (Fig. 2D), as well as comparing down-regulated genes in DNK group with the genes that their promoters were marked by Overlapped Peaks and Ctrl Specific Peaks. Venn plot identified 846 of the 1,163 up-regulated genes in DNK group, which consisting of receptor activity related Notch signaling component *notch2* are occupied with Overlapped Peaks and DN Specific H3K4me3 Peaks in the promoter regions (Fig. 2D, Figure 2-figure supplement 1C). These data suggest that endothelial specific inhibition of Brg1 results in increased H3K4me3 modification levels in the promoter region of genes, which in turn leads to up-regulation of genes expression, including *notch2*, in DN hearts.

### Endothelium-specific inhibition of Brg1 induces up-regulation of Notch signaling by increasing the level of H3K4me3 in the promoters

Since over-expression of DN-xBrg1 increases the levels of H3K4me3 modifications and mRNA expression of *notch2*, we then ask how Brg1 regulates *notch* receptor genes expression during heart regeneration. By performing RNA *in situ* analysis on frozen heart sections using either *notch1a*, *notch 1b*, *notch2*, or *notch 3* probes, we found that inhibition of Brg1 in endothelial cells (DN) resulted in slight up-regulation in sham-operated hearts, but had strong induction of *notch1a*, *notch1b*, *notch2*, and *notch3* in injured hearts at 7 dpa compared with control sibling hearts (Ctrl) (Fig. 3A). Furthermore, RNAscope *in situ* hybridization showed that *notch1b* overlapped with *kdrl*-positive endothelial cells but rarely with *tcf21*-positive epicardial cells in Ctrl hearts (Figure 3-figure supplement 1A, C) and DN hearts (Figure 3-figure supplement 1B, D). In addition, qRT-PCR of FACS-sorted *kdrl*:eGFP-positive endothelial cells from CtrlK and DNK hearts at 7 dpa showed that, compared with CtrlK group, the expression levels of *notch1a*, *notch1b*, *notch2*, and *notch3*, as well as Notch ligands *dll4*, significantly increased in DNK group (Fig. 3B) in the presence of 4-HT. Together, these data suggest an inhibitory effect of Brg1 on the expression of *notch* genes during heart regeneration.

**Figure 3.**
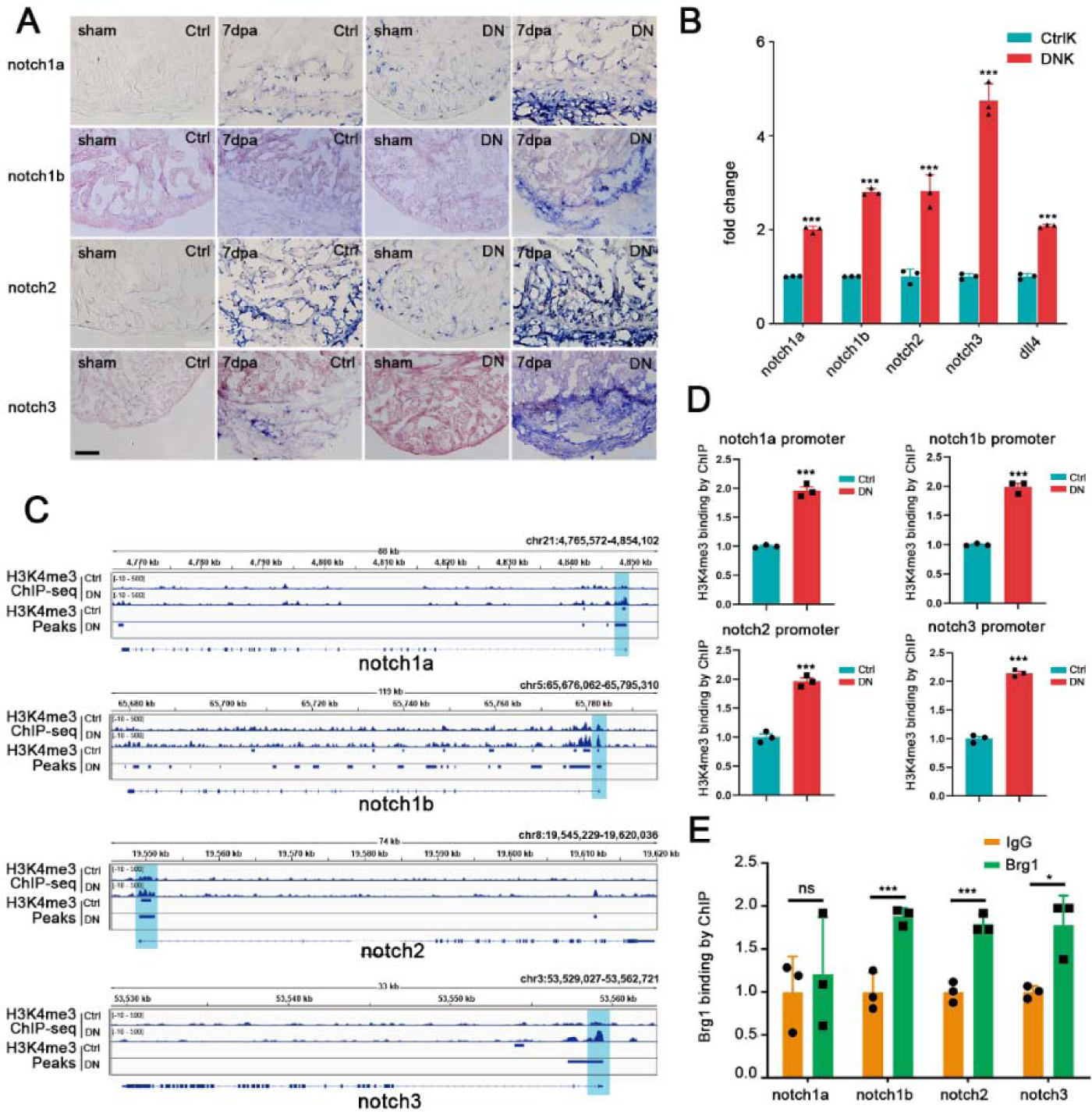
Endothelium-specific inhibition of Brg1 induces abnormal up-regulation of Notch signaling *via* the increased levels of H3K4me3 in their promoters. **(A)** Representative images of RNA *in situ* hybridization with *notch1a*, *notch1b*, *notch2*, and *notch3* probes on frozen sections of sham-operated Ctrl hearts, injured Ctrl hearts, sham-operated DN hearts, and injured DN hearts at 7 dpa. Scale bar, 100 μm. **(B)** Quantitative RT-PCR analysis showing that the expression of Notch receptors and ligand in FACS-sorted *kdrl*-eGFP positive endothelial cells from the DNK group is higher than those from the CtrlK group. Data represent one of three independent experiments, n=3 technical replicates for each group. Data are mean fold changes after normalization to GAPDH and expressed as the mean ± s.e.m., ***p <0.005, unpaired *t*-test. **(C)** H3K4me3 ChIP-seq showing the traces and peak intervals of representative genomic loci from Ctrl and DN hearts. Subtraction of normalized read coverage of H3K4me3 signals is shown in the displayed genomic windows. H3K4me3 peaks in both Ctrl and DN groups are shown as bars. Putative promoter regions are indicated in blue color. **(D)** Anti-H3K4me3 ChIP and quantitative PCR in Ctrl and DN hearts at 7dpa (primers designed from Notch receptor genomic regions: *notch1a*, –171/+3 bp; *notch1b*, – 41/+58 bp; *notch2*, –263/–115 bp; *notch3*, +394/+504 bp; ATG site designed as +1 bp). Data represent one of three independent experiments, n=3 technical replicates for each group. Data are the mean fold changes ± s.e.m.; ***p <0.005, unpaired *t*-test. **(E)** Anti-Brg1 ChIP and quantitative PCR in wild-type hearts at 3 dpa. Note the enrichment of Brg1 binding to Notch receptor promoters (*notch1b*, *notch2*, and *notch3*) but not to the *notch1a* promoter. Data represent one of two independent experiments, n=3 technical replicates for each group. Data are the mean fold change ± s.e.m.; *p <0.05, ***p <0.005; unpaired *t*-test. **Figure 3-source data 1**. Source data for Figure 3B, D, E.

We then investigated how Brg1 regulated Notch receptor genes. Genome-wide H3K4me3 ChIP-seq data showed that the H3K4me3 levels and peaks were increased in the promoters of *notch1a*, *notch1b*, *notch2*, and *notch3* genomic loci in DN group compared with those in Ctrl group (Fig. 3C). Particularly, the promoter regions of *notch1a*, *notch1b* and *notch2* occupied with the Overlapped Peaks and the peaks signals were stronger in the DN group compared with Ctrl group; and the promoter region of *notch3* had a H3K4me3 peak in DN group that was not in Ctrl group (Fig. 3C). We then used ChIP-qPCR to further confirm the levels of H3K4me3 modification in each of the Notch promoter regions. ChIP with H3K4me3 antibody and quantitative PCR (ChIP-qPCR) showed that the levels of H3K4me3 of all four Notch promoter regions were higher in DN hearts than in Ctrl hearts at 7 dpa in the presence of 5 μM 4-HT for 3 days before surgery (Fig. 3D), which was consistent with the elevated expression levels of these genes upon endothelial Brg1 inhibition (Fig. 3A, B). Furthermore, ChIP-qPCR with Brg1 antibody showed that Brg1 bound to the promoter regions of *notch1b*, *notch2*, and *notch3* but not *notch1a* (Fig. 3E), suggesting that Brg1 is involved in regulating the H3K4me3 modifications in the Notch promoters.

### Abnormally-activated Notch signaling is responsible for the reduced cardiomyocyte proliferation in DN-xBrg1 hearts

Since a previous study has shown that hyperactivation of Notch signaling impairs cardiomyocyte proliferation and heart regeneration (Zhao et al., 2014), we suspected that abnormally-activated Notch signaling might contribute to defects of cardiomyocyte proliferation and regeneration in the endothelium-specific DN-xBrg1 hearts. We generated Tg(*ubi*:loxp-DsRed-STOP-loxp-NICD; *kdrl*:CreER) transgenic fish line to carry out tamoxifen-inducible over-expression of NICD (zebrafish *notch1b* intracellular domain) that specifically activated Notch signaling in endothelial cells. Compared with control hearts at 7 dpa, we found that hyperactivation of Notch signaling in endothelial cells decreased the numbers of PCNA^+^/Mef2C^+^ proliferating cardiomyocytes (Fig. 4A-C), which was consistent with the previous report (Zhao et al., 2014). We then asked whether simultaneous knockdown of Notch receptors could rescue the numbers of proliferating cardiomyocytes in DN-xBrg1 mutant hearts. As described above, control and DN-xBrg1 zebrafish were infused with 5 μM 4-hydroxytamoxifen (4-HT) for 3 days before surgery, and nanoparticle-encapsulated *notch1a*, *notch1b*, *notch2*, *notch3* or control siRNA was, respectively, injected every day after surgery until the hearts were harvested at 7 dpa. With control siRNA injection, we found that the PCNA^+^/Mef2C^+^ proliferating cardiomyocytes were fewer in DN hearts than in Ctrl hearts (Fig. 4D, E, J). Interestingly, either *notch1a*, *notch1b*, *notch2*, or *notch3* siRNA was able to partially rescue the numbers of PCNA^+^/Mef2C^+^ proliferating cardiomyocytes in DN zebrafish hearts at 7 dpa, but was unable to return them to the control level (Fig. 4D-J), suggesting that hyperactivation of Notch signaling contributes to defects of myocardial proliferation in DN mutant hearts. In addition, we also chose two chemical inhibitors DAPT and MK-0752 to interfere with Notch signaling. Compared with DN hearts injected with control DMSO (Fig. 4L), we found more PCNA^+^/Mef2C^+^ proliferating cardiomyocytes in the DN hearts injected with either of the Notch inhibitors (Fig. 4M, N), but fewer than those in Ctrl hearts injected with DMSO (Fig. 4K, O). Thus, these results suggest that abnormally-activated Notch signaling is partially responsible for the cardiomyocyte-proliferation defects in the DN mutant heart. Together, our data suggest that injury-induced endothelial Brg1 negatively regulates the level of H3K4me3 in the promoter regions of *notch1b*, *notch2*, and *notch3*, and thus prevents the over-activation of notch signaling during heart regeneration. When this suppression is released, such as in DN hearts, the level of H3K4me3 modifications in the Notch promoter regions is abnormally up-regulated, resulting in the over-activation of Notch signaling and thus inhibiting regeneration.

**Figure 4.**
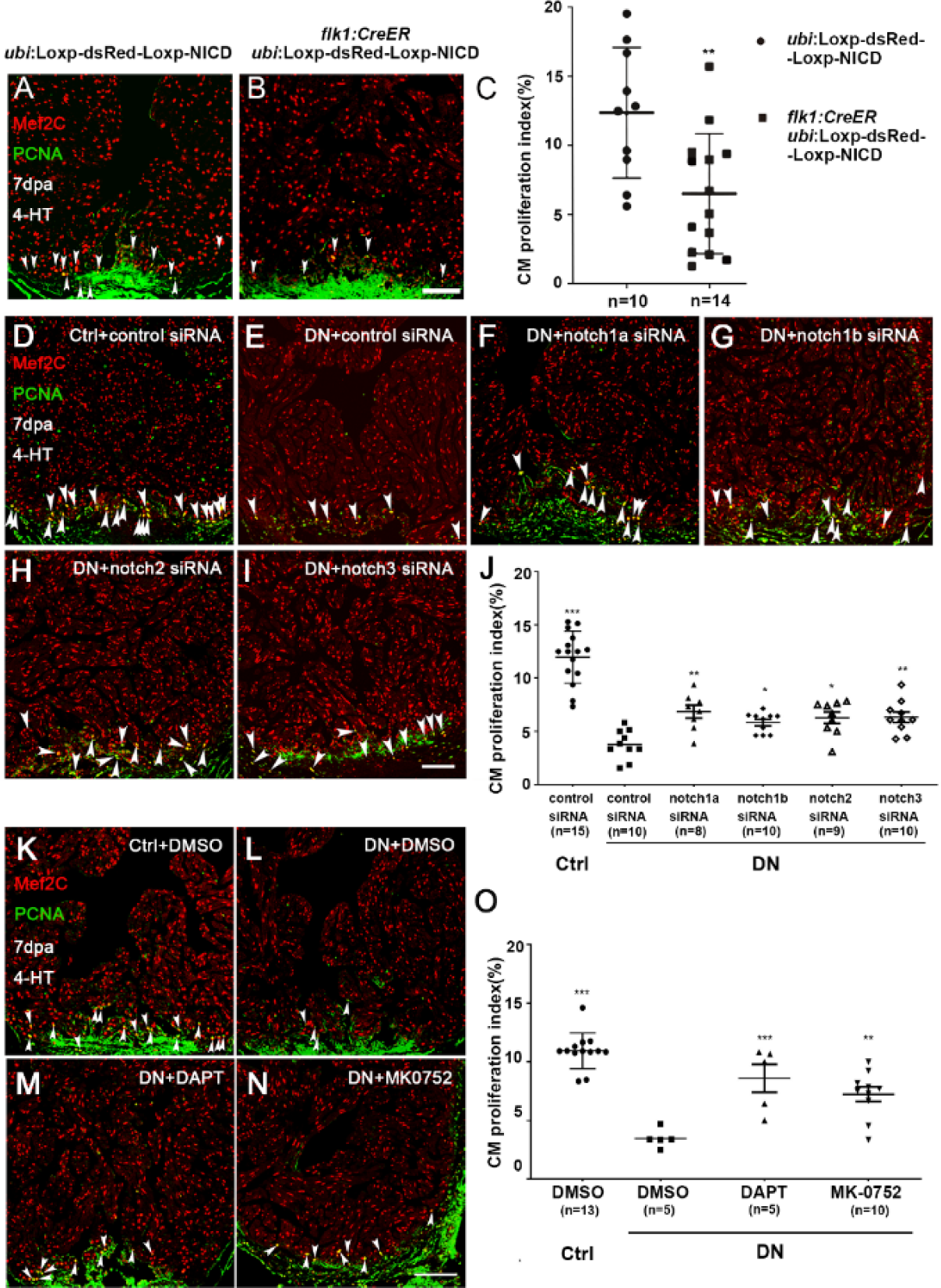
Endothelium-specific expression of NICD or DN-xBrg1 decreases cardiomyocyte proliferation that is partially rescued by inhibition of Notch signaling. **(A, B)** Immunostaining showing that Mef2C^+^ and PCNA^+^ proliferating cardiomyocytes of control (A) and endothelial NICD-overexpressing heart sections (B) at 7 dpa after 4-HT induction. (**C**) Statistics of panels **A** and **B** (data are the mean fold-change ± s.e.m.; **p <0.01, unpaired *t*-test). **(D**-**I)** Representative images of immunostaining showing that, compared with control siRNA treatment (D), PCNA^+^/Mef2C^+^ proliferating cardiomyocytes decreased at 7 dpa in DN-xBrg1 hearts (DN) (E), which were partially rescued by either *notch1a* (F), *notch1b* (G), *notch2* (H), or *notch3* (I) siRNA treatment in the presence of 4-HT. Scale bar, 100 μm. (**J**) Statistics of panels D-I (data are the mean ± s.e.m.; *p <0.05; **p <0.01; ***p <0.005; one-way analysis of variance followed by Dunnett’s multiple comparison test). **(K**-**N)** Representative images of immunostaining at 7 dpa showing that, compared with DMSO treatment (**K**), PCNA^+^/Mef2C^+^ proliferating cardiomyocytes in DN mutant hearts decreased (L), which were partially rescued by either DAPT (M) or MK-0752 treatment (N) in the presence of 4-HT. Scale bar, 100 μm. (**O**) Statistics of panels K-N (data are the mean ± s.e.m.; ***p<0.005; one-way analysis of variance followed by Dunnett’s multiple comparison test). N number shown here (C, G, O) indicate biological replicate. **Figure 4-source data 1.** Source images for Figure 4A-B, D-I, K-N. **Figure 4-source data 2.** Source data for Figure 4C, J, O.

### Brg1 interacts with Kdm7aa to fine-tune Notch signaling

We then asked how Brg1 negatively regulated H3K4me3 modifications in the promoter regions and had its function in regulating Notch receptor gene expression. It has been reported that Brg1 and histone demethylase (lysine demethylases, KDMs) jointly regulated gene expression in other organs (Li et al., 2019; Liu et al., 2019; Zhang et al., 2019). To determine whether KDMs were involved in the regulation of the levels of H3K4me3 by Brg1, we first examined the expression pattern of KDMs during zebrafish heart regeneration. RT-PCR data revealed that *kdm7aa* had the strongest expression while *kdm1a*, *kdm3b*, *kdm5bb*, *kdm6a*, *kdm6ba*, and *kdm6bb*, but not *kdm7ab* and *kdm8*, were weakly expressed in injured hearts at 2 dpa (Fig. 5A).

**Figure 5.**
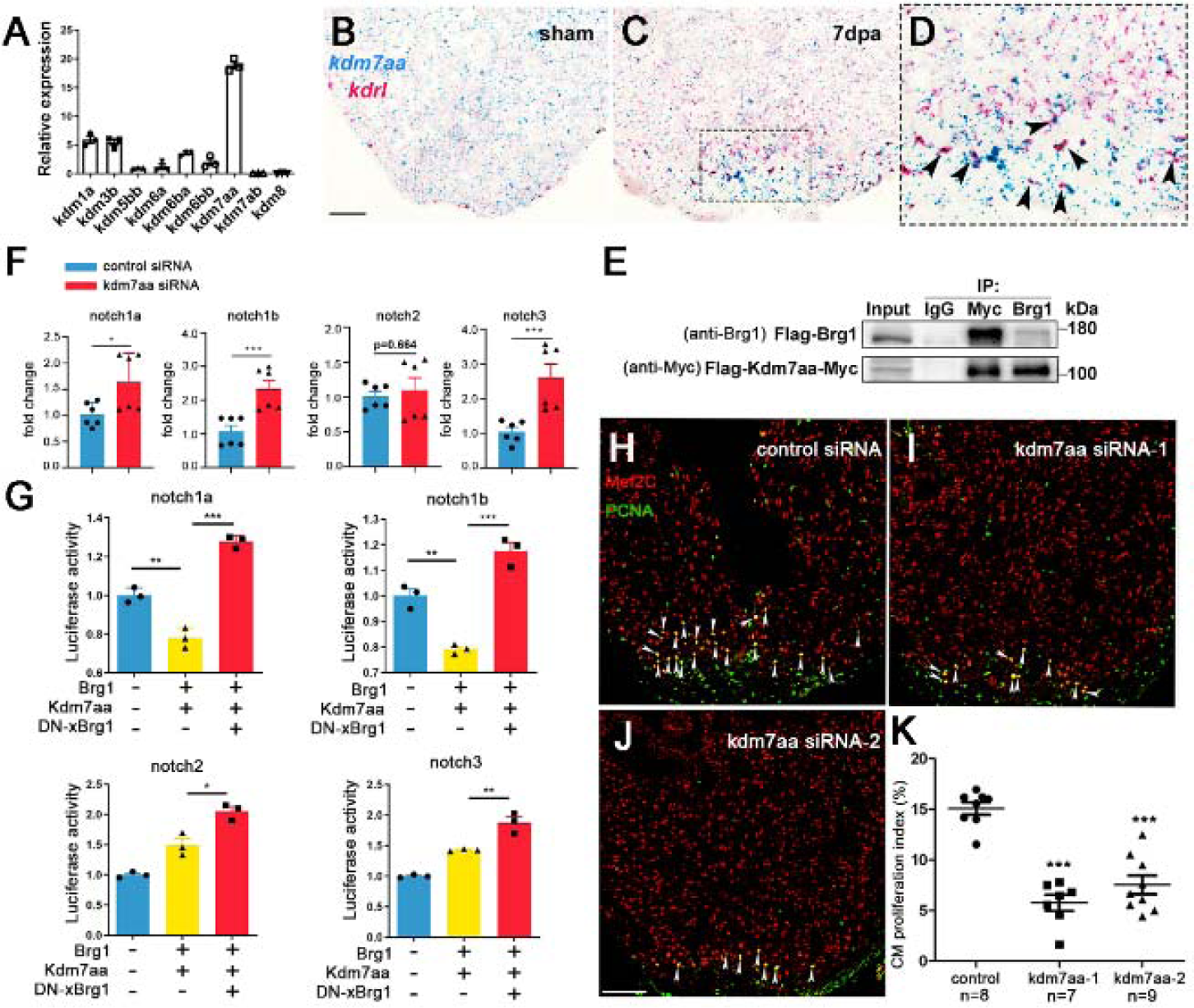
Endothelial Brg1 regulates Notch receptor expression and myocardial proliferation *via* interaction with Kdm7aa. **(A)** Quantitative RT-PCR of KDM genes expression, normalized by GAPDH, showing that *kdm7aa* has the highest expression level in injured zebrafish hearts at 2 dpa, n=3 technical replicates for each group. **(B-D)** Representative images of RNAscope *in situ* hybridization with *kdrl* and *kdm7aa* probes, showing that *kdm7aa* was expressed in sham (B) and injured hearts (C), and particularly overlaped with injury-induced *kdrl* expression in endothelial cells at 7 dpa (D) (scale bar, 100 μm) and high-magnification image of boxed region in D (arrowheads, double *kdrl*- and *kdm7aa*-positive endothelial cells). **(E)** Immunoprecipitation (IP) assays with either anti-Myc or anti-Brg1 antibody showing the interaction between Flag- Kdm7aa-Myc and Flag-Brg1 in 293T cells. Inputs used as loading controls and IgG as negative controls. **(F)** Quantitative RT-PCR analysis showing that the expression of *notch1a*, *notch1b*, *notch2* but not *notch3* from hearts at 7 dpa injected with encapsulated kdm7aa siRNA was higher compared with control siRNA group. Data represent one of two independent experiments, n=6 (2 biological replicates with 3 technical replicates per biological sample). Data are mean fold changes after normalization to GAPDH and expressed as the mean ± s.e.m., *p <0.05, ***p <0.005, unpaired *t*-test. **(G)** Luciferase reporter assays in 293T cells stably expressing the Notch promoter-luciferase reporter in the pGL4.26 vector. Expression plasmid clones containing Kdm7aa, Brg1, or DN-xBrg1 were co-transfected into cells stably expressing each Notch reporter. Data represent one of two independent experiments, n=3 technical replicates for each group, *p <0.05, **p <0.01, ***p <0.005, one-way analysis of variance followed by Bonferroni test. **(H**–**J)** Representative images of immunostaining showing that the numbers of Mef2C^+^/PCNA^+^ proliferating cardiomyocytes decreased in injured hearts at 7 dpa injected with either encapsulated *kdm7aa* siRNA-1 (I) or siRNA-2 (J) compared with control siRNA (H) (arrowheads, Mef2C^+^/PCNA^+^ proliferating cardiomyocytes; scale bar, 100 μm). (**K**) Statistics of panels H–K (n numbers indicated biological replicates, data are the mean ± s.e.m.; ***p<0.005; one-way analysis of variance followed by Dunnett’s multiple comparison test). **Figure 5-source data 1.** Source data for Figure 5A, F, G, K. **Figure 5-source data 2.** Raw Western Blot for Figure 5E and Source data for Figure 5H-J.

Kdm7aa has been shown to be responsible for histone demethylation at multiple sites, including H3K9, H3K27, H3K36, and H3K20 (Tsukada, Ishitani, & Nakayama, 2010). Interestingly, we also found that *kdm7aa* was induced and enriched in cardiac endothelial cells upon injury using RNAscope with *kdrl* and *kdm7aa* probes (Fig. 5B-D). Therefore, we further examined the interaction between Brg1 and Kdm7aa using immunoprecipitation (IP). Lysates of cells over-expressing both Flag-Kdm7aa-Myc and Flag-Brg1 were precipitated by either Myc or Brg1 antibody. Western blots revealed that IP with either Myc antibody (Myc-tagged Kdm7aa) or Brg1 antibody was able to pull down both Flag-tagged Brg1 (∼180 kD) and Myc-tagged Kdm7aa (∼100 kD), suggesting that Brg1 physically interacted with Kdm7aa (Fig, 5E). To examine whether Kdm7aa is involved in Brg1-regulated Notch receptor gene expression, we utilized nanoparticle-mediated gene-silencing (Diao et al., 2015; Xiao et al., 2018) to knockdown *kdm7aa*, RT-PCR results displayed that down-regulation of *kdm7aa* significantly up-regulated *notch1a*, *notch1b* and *notch3* expression (Fig. 5F). We also used the luciferase reporter system that were driven by *notch1a*-, *notch1b*-, *notch2*-, or *notch3* promoters, and made stable 293T cell lines expressing each of the luciferase reporters. Luciferase assays showed that over-expression of Brg1 and Kdm7aa decreased *notch1a* and *notch1b* reporter activity, while over-expression of DN-xBrg1 increased the activity of all four Notch reporters (Fig. 5G), suggesting a synergistic role of Brg1 and Kdm7aa in controlling the expression levels of Notch reporter genes. We finally set out to address whether *kdm7aa* was directly involved in regulating zebrafish heart regeneration. We found that knockdown of *kdm7aa* with two independent siRNAs decreased the numbers of PCNA^+^/Mef2C^+^ proliferating cardiomyocytes compared with control siRNA (Fig. 5H-K). Together, our data suggest that endothelial cell Brg1 interacts with Kdm7aa to maintain the normal activity of Notch gene promoters, and Kdm7aa modulates the level of H3K4me3 to fine-tune Notch gene expression during heart regeneration.

## Discussion

In this study, we showed that endothelial Brg1 was required for myocardial proliferation and regeneration in zebrafish; Brg1 interacted with Kdm7aa to fine-tune the level of H3K4me3 in the Notch receptor promoters and negatively regulated Notch gene expression during heart regeneration; and Kdm7aa was induced in cardiac endothelial cells and was required for myocardial proliferation. Therefore, our data reveal a new function of the endothelial Brg1-Kdm7aa axis in regulating Notch gene transcription, and the essential role of histone methylation *via* Kdm7aa in myocardial proliferation and regeneration in zebrafish.

Previous studies have shown that Brg1 plays an important role in oocyte genome activation, erythropoiesis, T-cell generation, erythropoiesis, vascular development, nerve development, heart development and regeneration (Bultman, Gebuhr, & Magnuson, 2005; Bultman et al., 2006; Chi et al., 2003; Eroglu, Wang, Tu, Sun, & Mivechi, 2006; Griffin, Brennan, & Magnuson, 2008; Hang et al., 2010; Seo, Richardson, & Kroll, 2005; Stankunas et al., 2008; Xiao et al., 2016). We here demonstrated that conditional inhibition of Brg1 function in endothelial cells including the endocardium led to increased cardiac fibrosis and compromised myocardial proliferation and regeneration. Either hypo- or hyper-activation of Notch signaling has been reported to impair cardiomyocyte proliferation and heart regeneration (Munch, Grivas, Gonzalez-Rajal, Torregrosa-Carrion, & de la Pompa, 2017; Raya et al., 2003; Zhao et al., 2019; Zhao et al., 2014), suggesting that the precise modulation of Notch family expression is essential for cardiac regeneration. Here, we present several layers of evidence to demonstrate that injury-induced Brg1 and Kdm7aa regulate Notch gene expression in cardiac endothelium and endocardium. Brg1 and Kdm7aa normally fine-tune the level of the histone marker H3K4me3 in the Notch gene promoters, thus preventing the abnormal hyperactivation of Notch receptors after injury. When Brg1 was inhibited in cardiac endothelial cells, the H3K4me3 level increased in the Notch promoter regions and Notch genes were abnormally over-expressed, leading to enhanced cardiac fibrosis and compromised myocardial proliferation and regeneration. Injury-induced expression of *brg1* and *kdm7aa* was evident in cardiac endothelial cells that was consistent with their function, which were further supported by our data on the physical interaction between Brg1 and Kdm7aa, and their function in regulating Notch promoter activities. Furthermore, either encapsulated siRNA knockdown of Notch receptors, or chemical Notch inhibitors, partially rescued the phenotype of myocardial proliferation in DN-xBrg1 hearts, further suggesting an important role of Brg1 in regulating Notch gene expression during heart regeneration. At the same time, how hyper-activated Notch signaling in cardiac endothelium and endocardium represses myocardial proliferation *via* endocardium-myocardium interaction warrants future investigations.

Chromatin remodeling has been reported to be essential for tissue/organ regeneration in urodeles and zebrafish (Martinez-Redondo & Izpisua Belmonte, 2020; Zhu et al., 2018). Brg1 is the major subunit of the SWI/SNF complex, and is also an important component of the trithorax group, both of which play essential roles in histone modification such as the histone markers H3K4me3 (active) and H3K27me3 (repressive). Although data on genome-wide histone acetylation and methylation during organ regeneration are still limited, recent studies suggest that a more open chromatin state is adopted during early fin, retina, and heart regeneration in zebrafish (Goldman et al., 2017; Stewart, Tsun, & Izpisua Belmonte, 2009; Wang et al., 2020). The level of the histone marker H3K4me3 is influenced and catalyzed by lysine methyltransferases of the MLL2 complex and KDMs. Although the MLL2 complex does not provide selective specificity in a particular organ or biological process, it is believed that ATP-dependent chromatin remodeling proteins such as Brg1 may specifically regulate the “bivalency” state of H3K4me3 and H3K27me3 (Harikumar & Meshorer, 2015). KDM7 has been reported to act as a dual KDM for histone silencing markers H3K9 and H3K27 in brain development and germ cell genome stability (Myers, Amendola, Lussi, & Salcini, 2018; Tsukada et al., 2010), but it is unknown whether it also works for the active histone marker H3K4. We found that *brg1* and *kdm7aa* co-expressed in cardiac endothelial cells upon injury in zebrafish, and they formed a protein complex and functioned synergistically to regulate Notch receptor gene promoters in mammalian cells. Inhibition of Brg1 function *via* DN-xBrg1 mutant proteins increased the Notch promoter activity, suggesting that DN-xBrg1 might replace and/or inhibit Kdm7aa function and so increased the level of H3K4me3. Furthermore, the data on nanoparticle-mediated kdm7aa siRNA knockdown supported its function in myocardial proliferation and regeneration. Thus, this work reveals an interesting mechanism on the selective modulation of H3K4me3 by Brg1 and Kdm7aa and their essential function in zebrafish heart regeneration.

## Acknowledgements

The authors thank Dr IC Bruce (Guest Professor at the Institute of Molecular Medicine, Peking University) for commenting and revising the manuscript, and members of Dr. Jing-Wei Xiong’s and Dr. Yan-Yi Huang’s laboratories for helpful discussions and technical assistance. The authors acknowledge Chenyang Geng, Yun Zhang, Jing Sun, and Yang Xu for technical assistance at the Beijing Advanced Innovation Center of Genomics and the High-Throughput Sequencing Center at Peking University; Fangjin Chen, Ting Fang, and Wenzhong Zhang for technical help at the Computing Platform of the Center for Life Sciences of Peking University; Li-Ying Du and Hong-Xia Lv for technical help at the Flow Cytometry Core at the National Center for Protein Sciences at Peking University. This work was supported by the National Key R&D Program of China (2019YFA0801602 and 2018YFA0800501) and grants from the National Natural Science Foundation of China (31701272, 31730061, 31430059 and 81870198) and. Chenglu Xiao was supported in part by postdoctoral fellowships from Peking University Boya Program and Peking-Tsinghua Center for Life Sciences.

## Competing Interests

The authors declare no competing interests.

## Materials and Methods

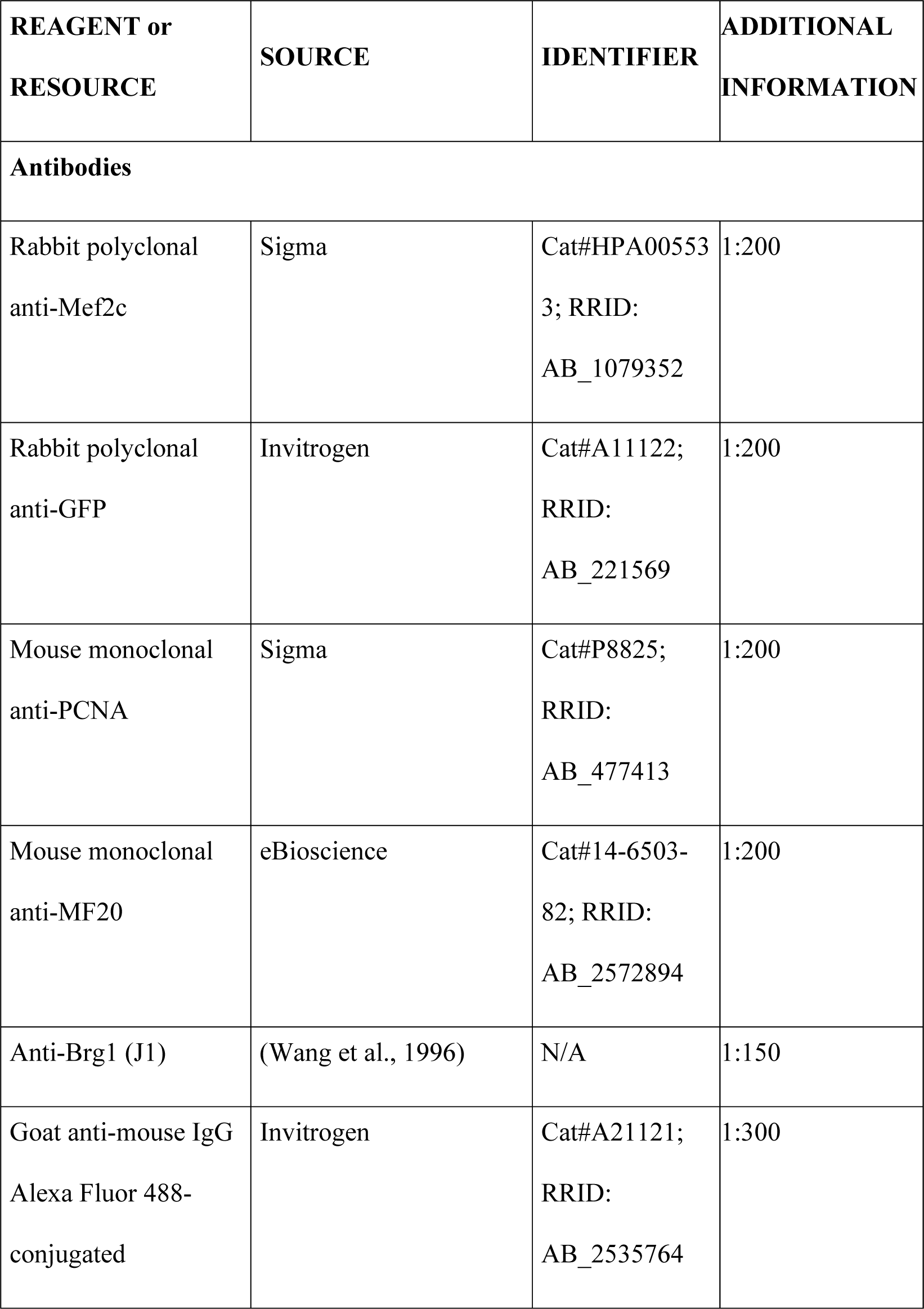
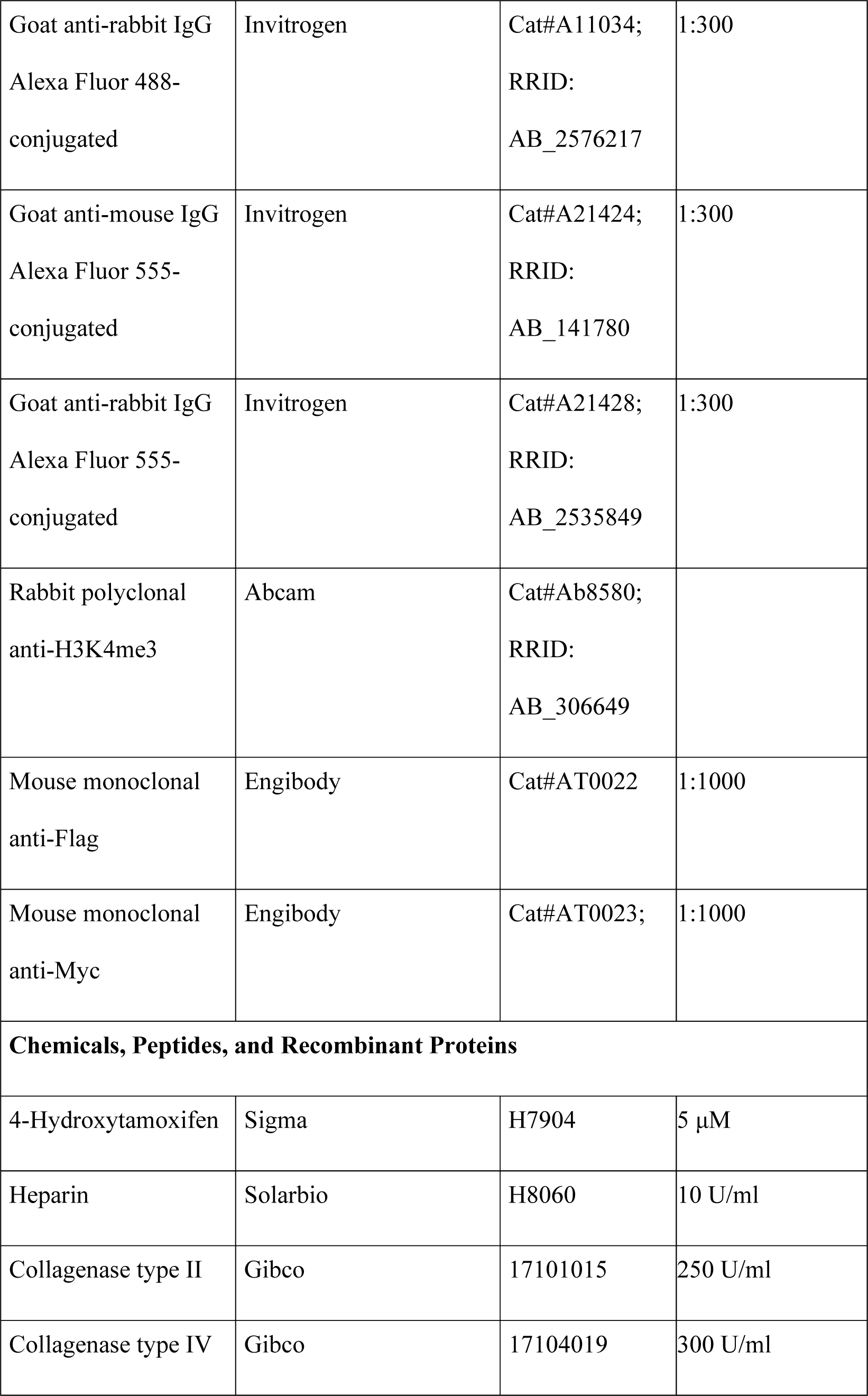
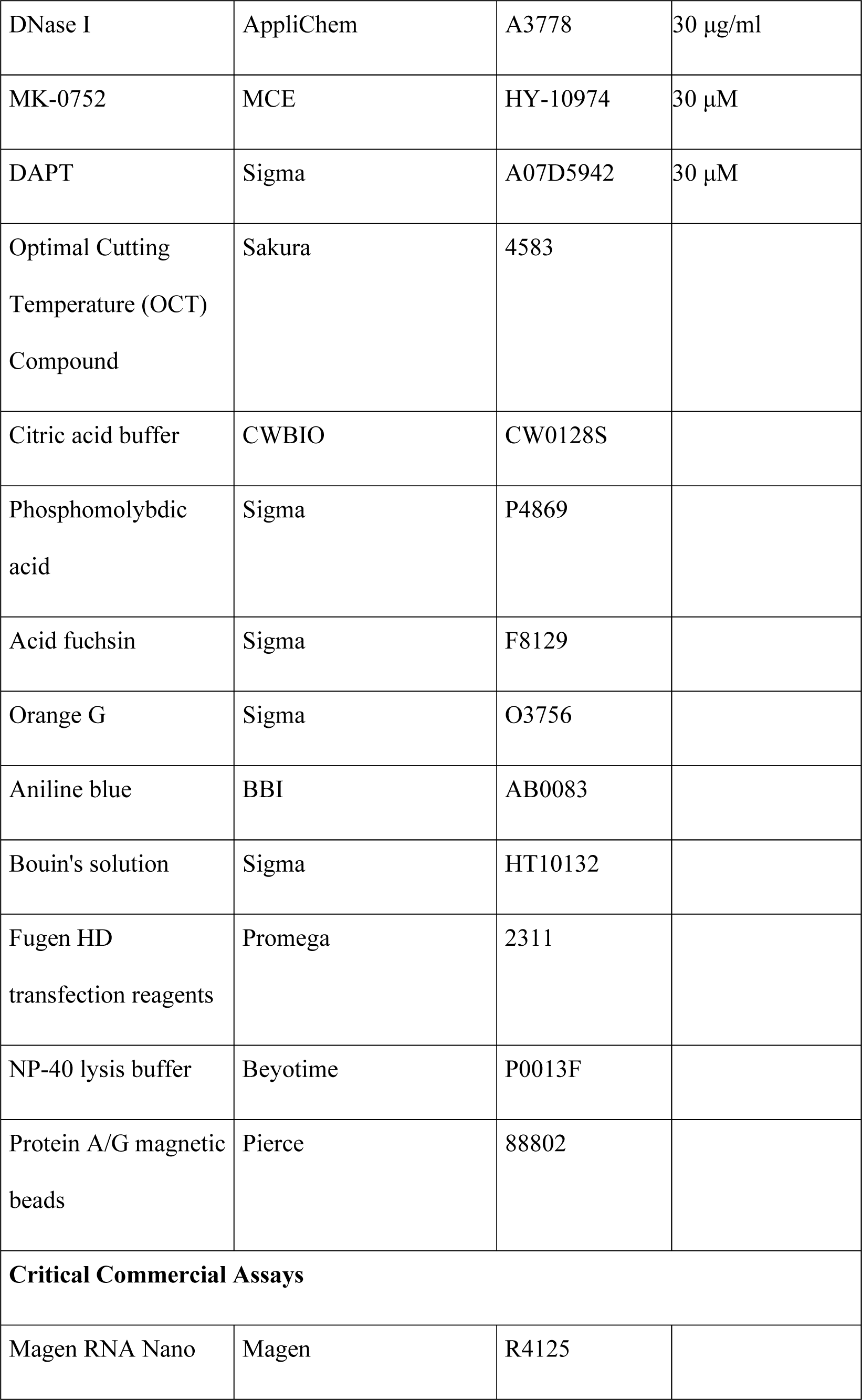
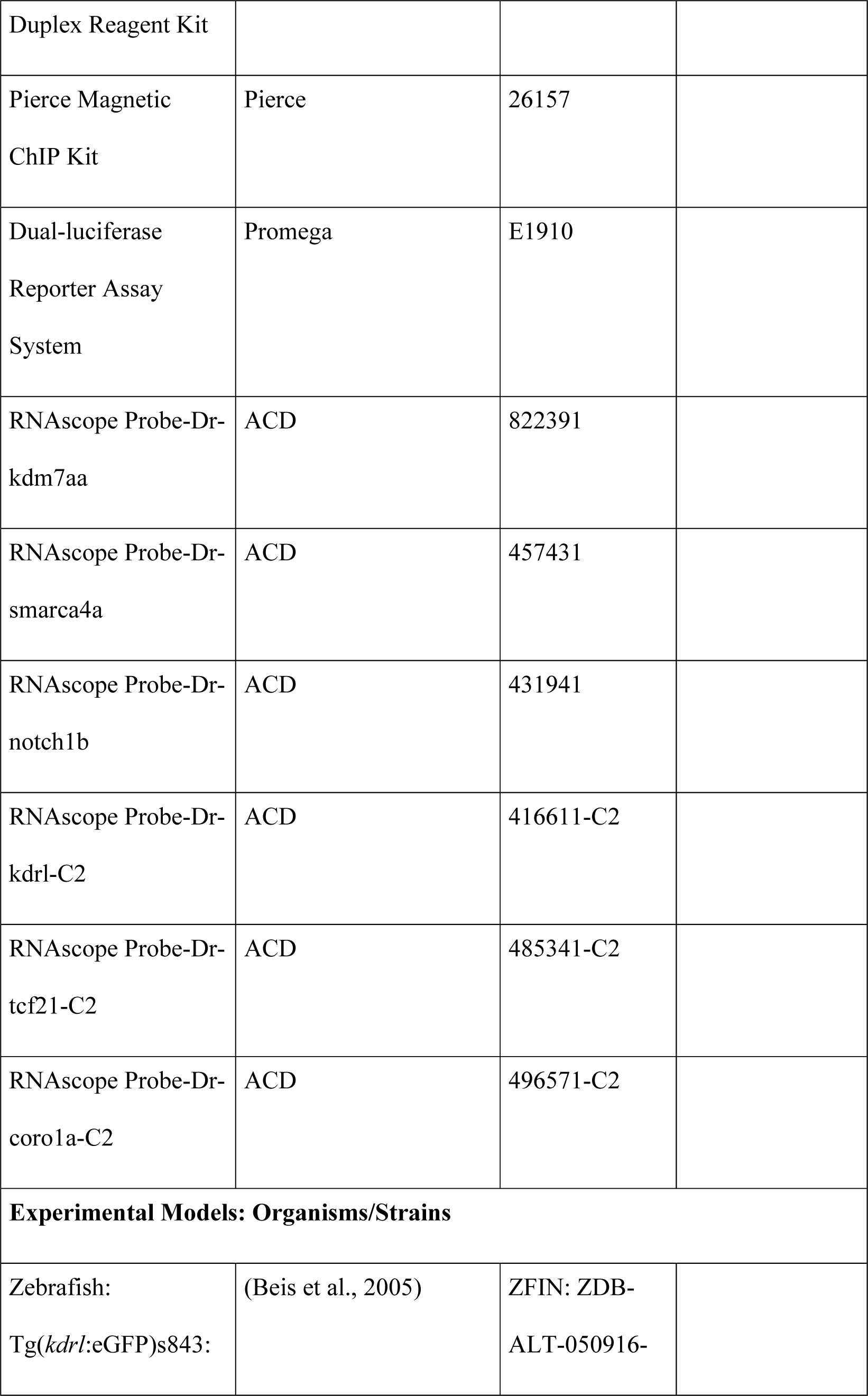
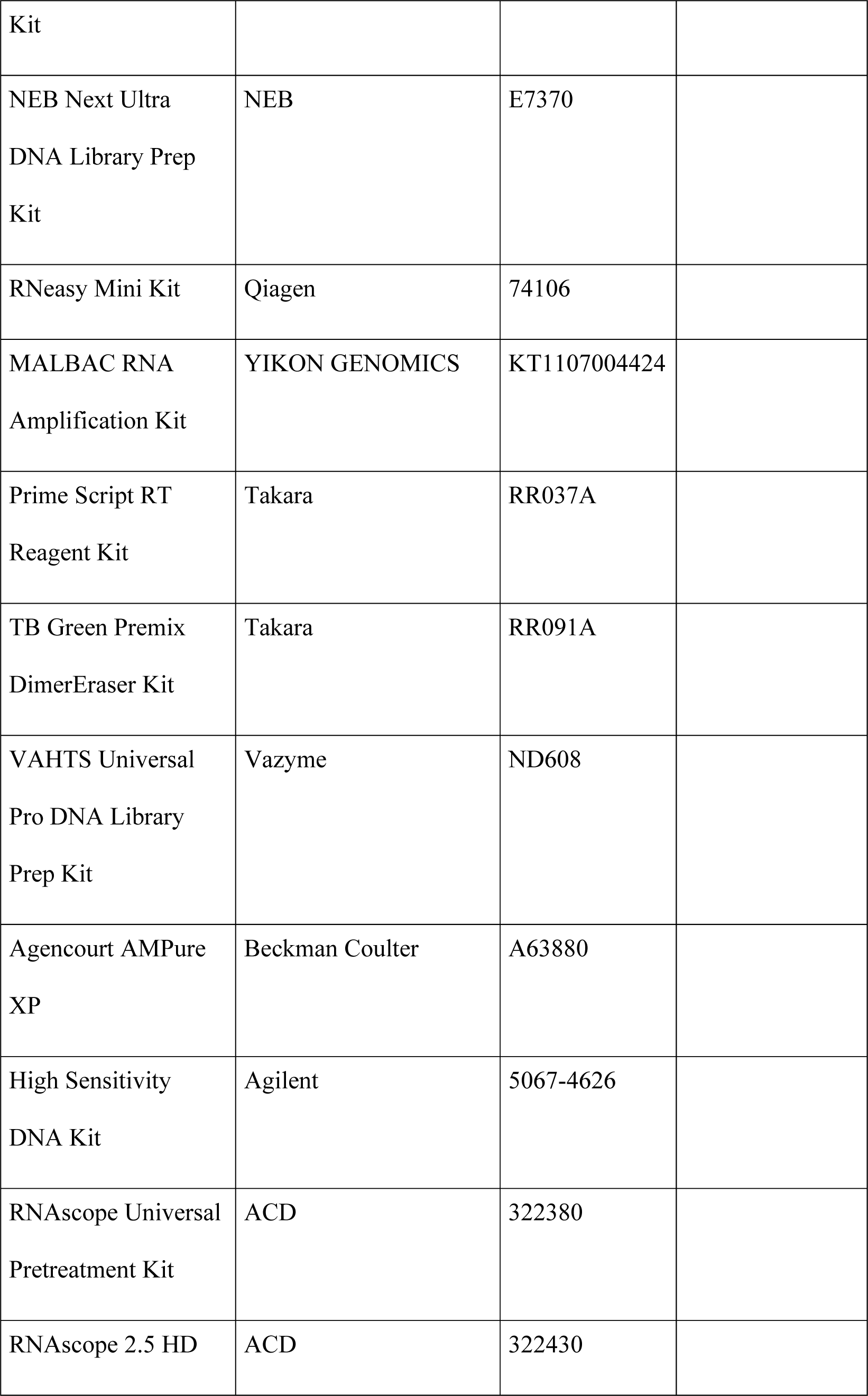
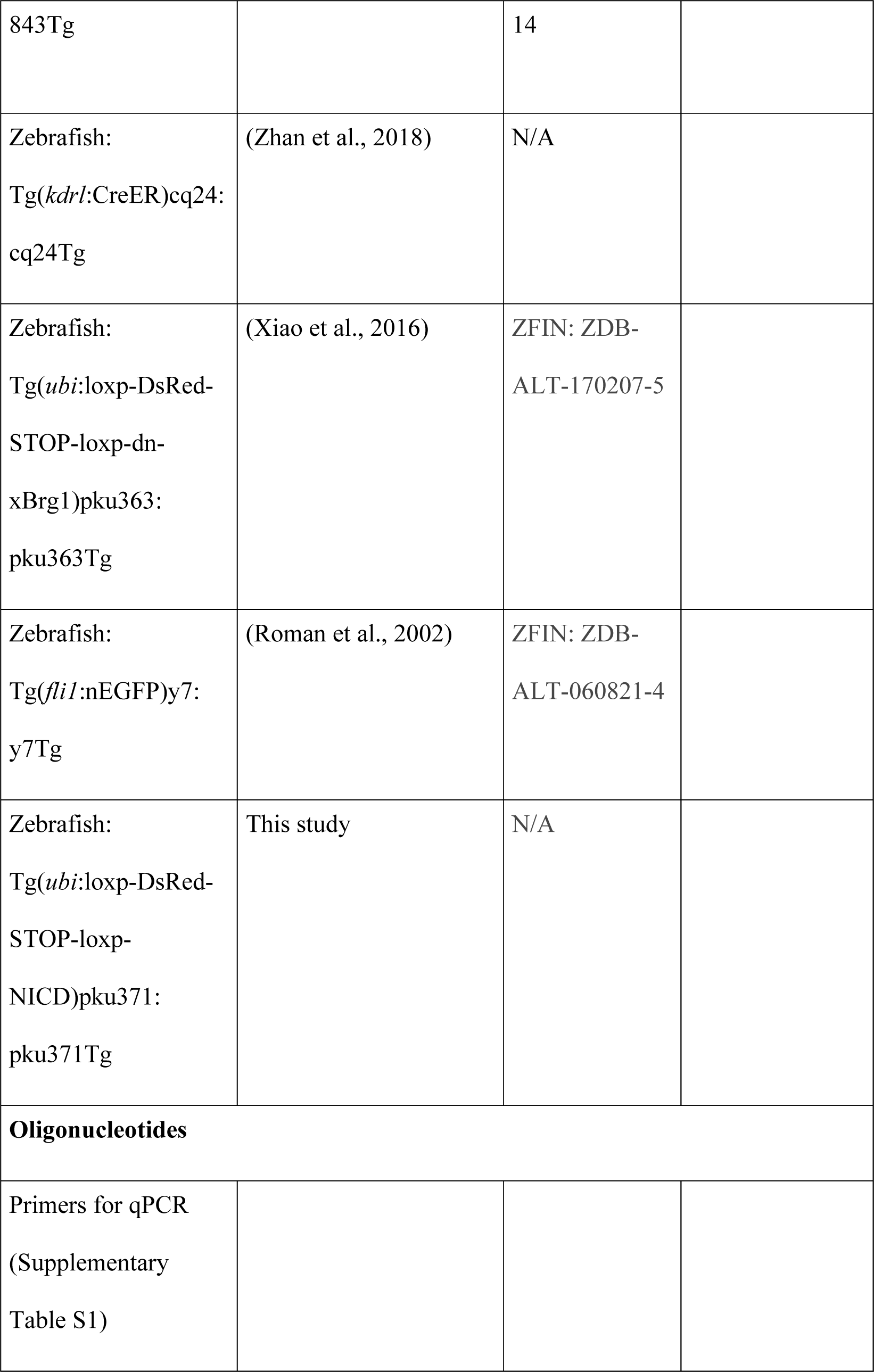
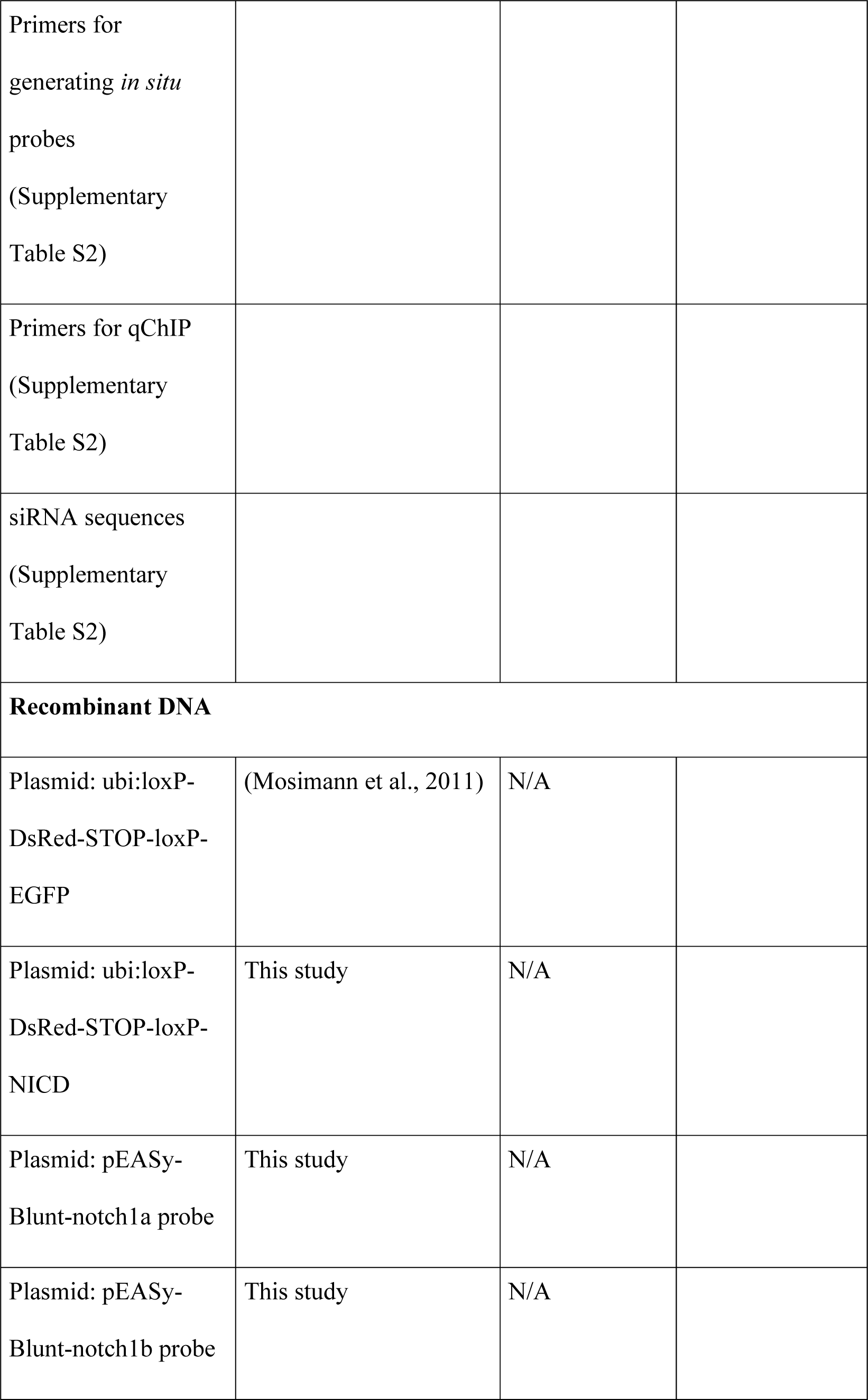
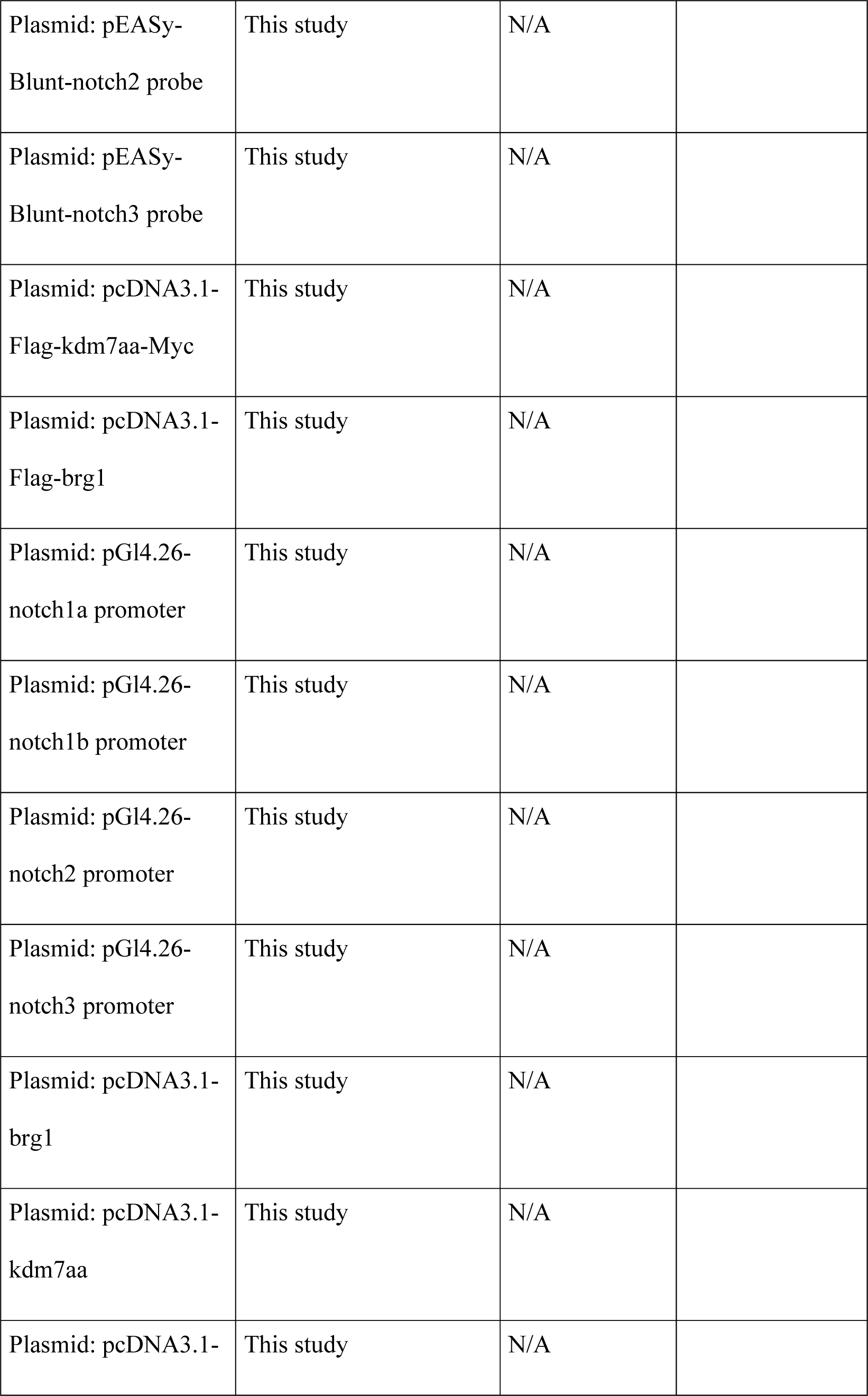
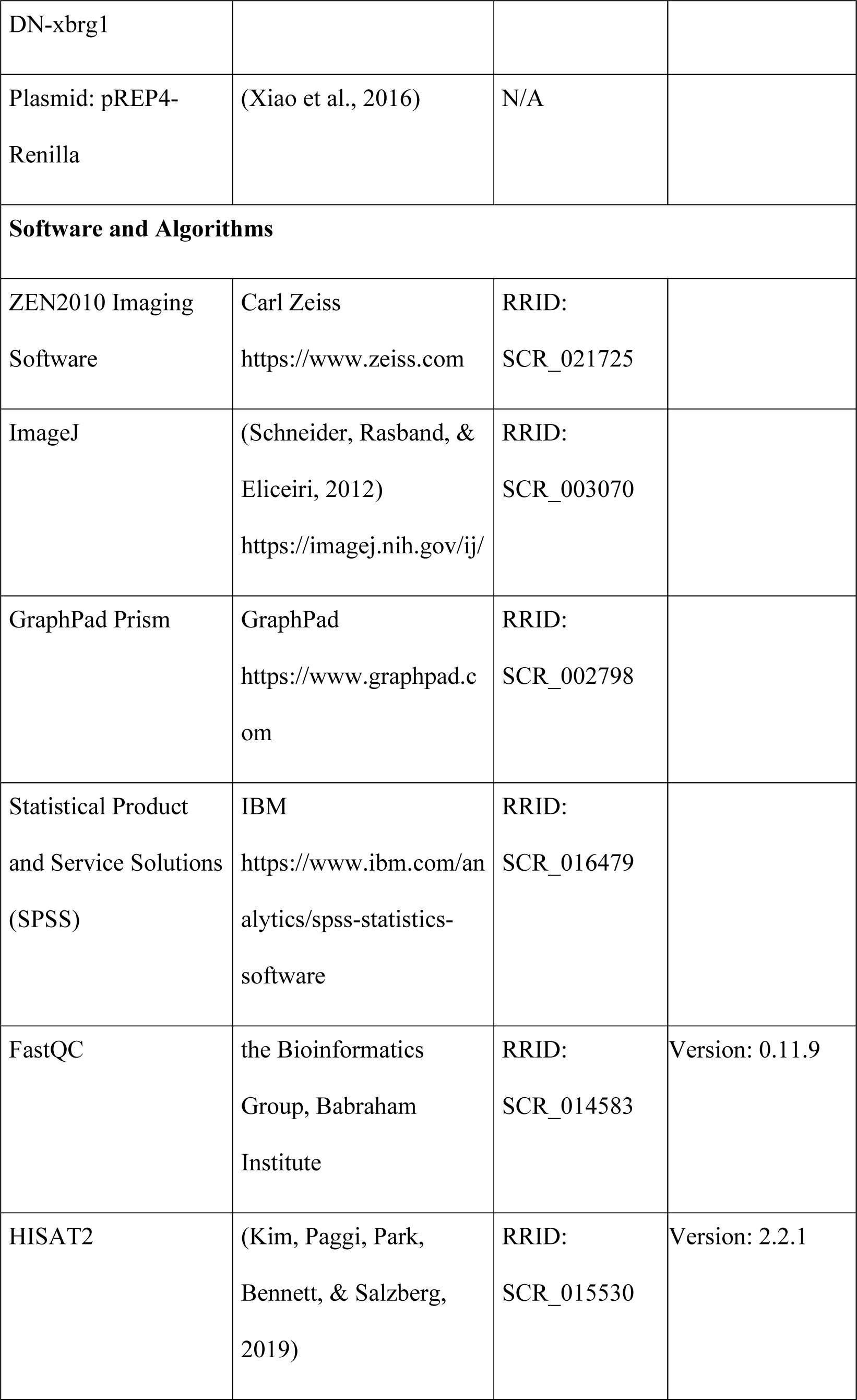
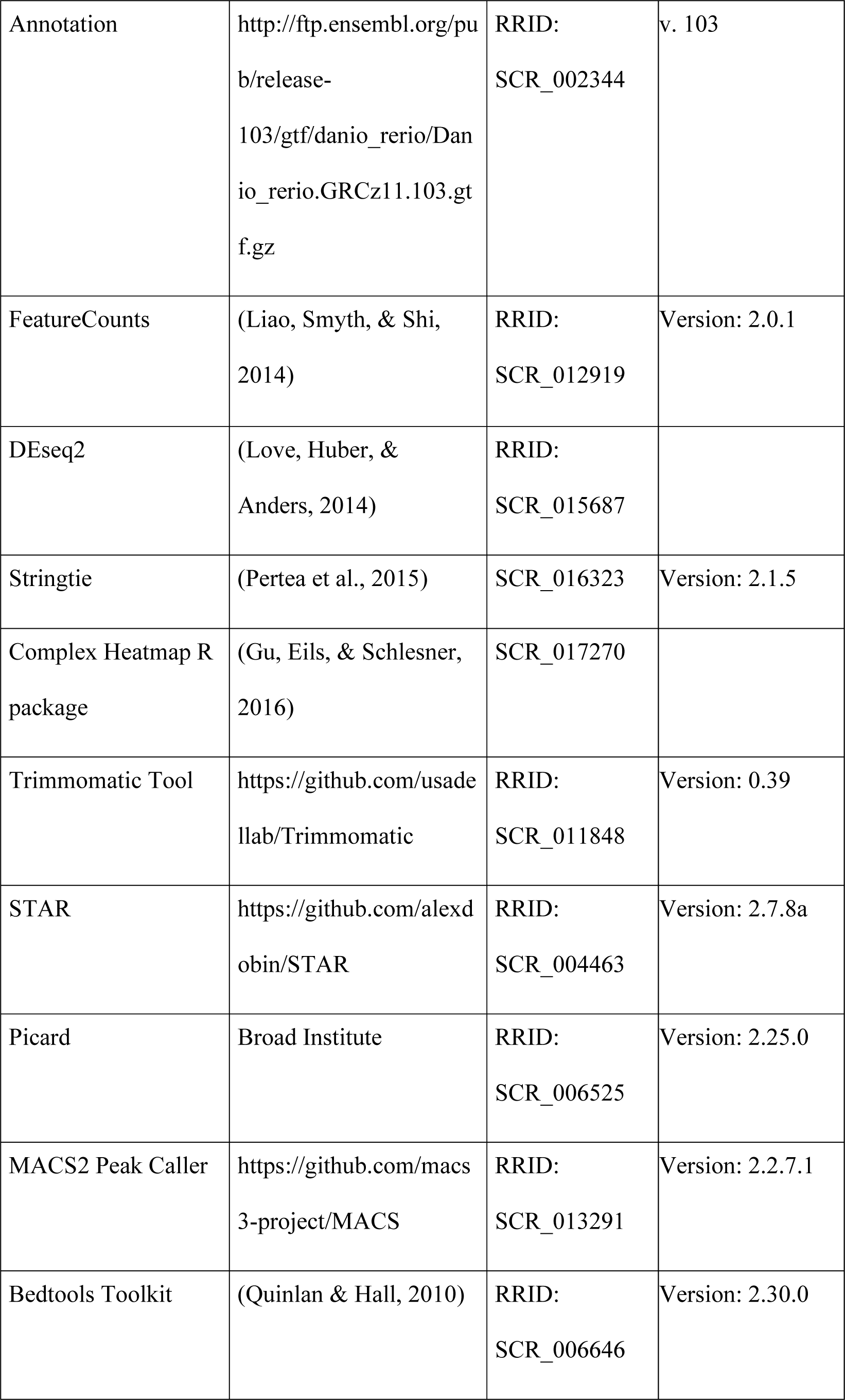
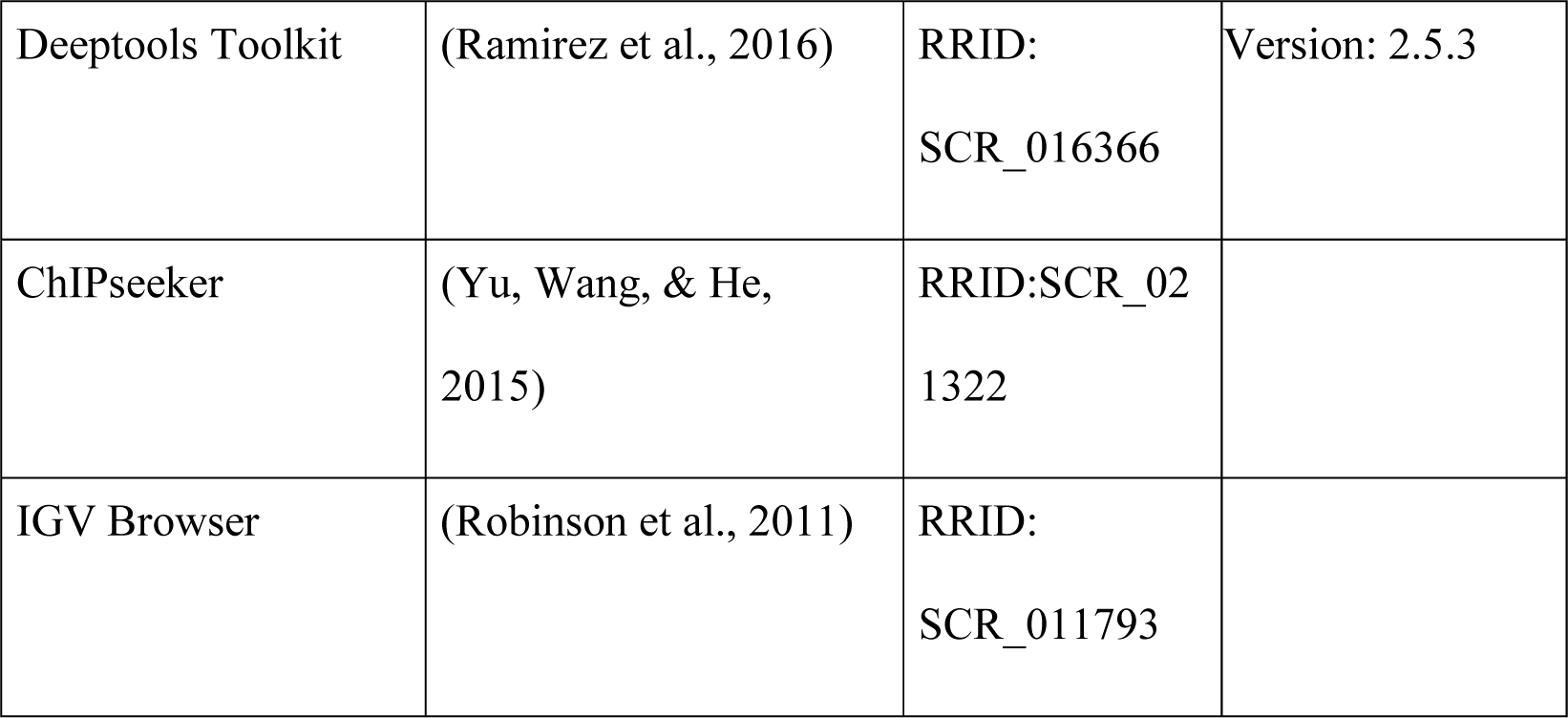

### Animal models

Male and female zebrafish were raised and handled according to a zebrafish protocol (IMM-XiongJW-3) approved by the Institutional Animal Care and Use Committee at Peking University, which is fully accredited by The Association for Assessment and Accreditation of Laboratory Animal Care International. Wild-type TU, Tg(*kdrl*:eGFP) (Beis et al., 2005), Tg(*kdrl*:CreER) (Zhan et al., 2018), Tg(*fli1*:nucEGFP) (Roman et al., 2002), and Tg(*ubi*:loxp-DsRed-STOP-loxp-DN-xBrg1) zebrafish (Xiao et al., 2016) were maintained at 28°C at a density of 4 fish per liter. Adult zebrafish were anesthetized in standard E3 medium containing 0.4% tricaine (ethyl 3-aminobenzoate methanesulfonate salt; Sigma-Aldrich) before ventricular resection as described previously (Xiao et al., 2016). Animals were randomized into groups for each experiment.

### Construction of Tg(*ubi*:loxp-dsRed-loxp-NICD) transgenic zebrafish line

To generate the Tg(*ubi*:loxp-DsRed-STOP-loxp-NICD) zebrafish line that over-express NICD, an homologous recombination reaction was conducted with ubi:loxP-DsRed-STOP-loxP-EGFP plasmid (kindly provided by Dr. C Geoffrey Burns at Harvard Medical School) (Mosimann et al., 2011) by replacing EGFP with zebrafish notch1b-NICD cDNA. This Tol2-NICD plasmid was made and injected into one-cell stage wild-type embryos together with Tol2 transposase mRNA as described previously (Kawakami et al., 2004). Heterozygous transgenic zebrafish were raised and genotyped for all experiments.

### 4-hydroxytamoxifen (4-HT) treatment

We generated Tg(*ubi*:loxp-DsRed-STOP-loxp-DN-xBrg1; *kdrl*:CreER) mutant (DN) and Tg(*ubi*:loxp-DsRed-STOP-loxp-DN-xBrg1) control sibling (Ctrl) adult zebrafish by crossing Tg(*ubi*:loxp-DsRed-STOP-loxp-DN-xBrg1) with Tg(*kdrl*:CreER) zebrafish. To induce Cre recombination, adult DN mutant and Ctrl sibling zebrafish were bathed for 24 h in the presence of 5 μM 4-HT (H7904; Sigma) made from a 10 mM stock solution dissolved in 100% ethanol at room temperature. These zebrafish were treated with 4-HT at a density of 3-4 zebrafish per 150 ml system water. Ventricular resection was performed 3 days after 4-HT treatment. Transgenic zebrafish were confirmed by PCR-based genotyping and were randomly selected for all experiments.

### Ventricular resection in adult zebrafish

The ventricular resection was performed according to a well-established procedure (Han et al., 2014; Poss, Wilson, & Keating, 2002; Xiao et al., 2018). Briefly, zebrafish were anaesthetized with 0.4% tricaine and placed in the groove of a sponge. The pericardial sac was exposed by removing surface scales and a small piece of skin and the ventricle apex was gently pulled up and removed with Vannas scissors. The zebrafish was quickly placed back into a system water tank, and water was puffed over the gills with a plastic pipette until it breathed and swam regularly. The surface opening sealed automatically within a few days.

### Fluorescence-activated cell sorting (FACS) of cardiac endothelial cells

Cardiac endothelial cells from Tg(*ubi*:loxp-DsRed-STOP-loxp-DN-xBrg1; *kdrl*:eGFP) control (CtrlK) and Tg(*ubi*:loxp-DsRed-STOP-loxp-DN-xBrg1; *kdrl*:CreER; *kdrl*:eGFP) mutant (DNK) ventricles at 7 dpa with 4-HT treatment were isolated according to an established protocol (Patra et al., 2017). Briefly, ∼15 adult zebrafish hearts were isolated and washed in cold PBS with 10 U/ml heparin (H8060; Solarbio). After the atrium and bulbus were removed, the ventricles were carefully cut into small pieces using forceps and collected into 1.5-ml centrifuge tubes containing cold PBS with 5 mM glucose. The sliced tissue was then transferred to a glass tube along with a magnetic stir bar and 1.5 ml digestion buffer in Dulbecco’s modified Eagle’s medium containing collagenase type II (250 U/ml) (17101015; Gibco), collagenase type IV (300 U/ml) (17104019; Gibco), and DNase I (30 μg/ml) (A3778; AppliChem). The tube was then transferred to a 32°C water bath with stirring and incubated for 1 min. After incubation, the tube was removed from the water bath and left at room temperature until the tissue settled on the bottom. The supernatant was discarded to remove blood cells, followed by washing once with cold PBS. This was followed by a series of digestion steps with 1.5 ml digestion buffer. Each step consisted of 10 min of digestion followed by 3 min of sedimentation. The supernatants were collected in a 15-ml falcon tube containing 2 ml ice-cold PBS. The cell suspensions were centrifuged at 300 g for 5 min at 4°C, and the cell pellets were gently re-suspended in 1 ml PBS kept on ice for FACS. Cardiac endothelial cells were sorted through the GFP channel and were collected into a tube containing 0.5 ml PBS with 10% FBS. The cells were centrifuged at 500 g for 5 min at 4°C, and the cell pellets were collected and kept on ice ready for RNA isolation.

### RNA-seq of cardiac endothelial cells

The RNA of heart endothelial cells from CtrlK sibling and DNK mutant ventricles at 7 dpa was purified using a Magen RNA Nano Kit (R4125; Magen). 30 ng of total RNA was used for next-generation library preparation under the guidelines of the NEBNext Ultra DNA Library Prep Kit for Illumina (E7370; NEB). The libraries were loaded for 2 × 150 bp pair-end sequencing using Illumina Hiseq 2500. Raw reads were pre-processed and quality controlled with FastQC (Version: 0.11.9). Reads for each library were mapped using HISAT2 (Version: 2.2.1) (Kim et al., 2019) against the zebrafish reference genome assembly GRCz11 with default parameters. Uniquely mapped reads were extracted to calculate the read counts of each gene, using the matching gene annotation (v. 103) from Ensembl with featureCounts (Version: 2.0.1). Genes were further filtered, and those with low expression in all samples (FPKM < 0.5 in all samples) were removed from differential gene expression analysis. Differential analysis was conducted with DEseq2 (Love et al., 2014). Genes with an adjusted P-value <0.05 were taken as significantly differentially expressed genes in the DNK condition compared with CtrlK. FPKM values were calculated with Stringtie (Version: 2.1.5) and Normalized Z-score values were used to draw heatmaps using the ComplexHeatmap R package (Gu et al., 2016). Sequencing data have been deposited in GEO under accession code GSE200936, https://www.ncbi.nlm.nih.gov/geo/query/acc.cgi?acc=GSE200936.

### ChIP-seq

ChIP-seq libraries were prepared using the VAHTS Universal Pro DNA Library Prep Kit (ND608; Vazyme) for Illumina. 5 nanogram of DNA was used as starting material for input and IP samples. Libraries were amplified using 13 cycles on the thermocycler. Post amplification libraries were size selected at 250-450bp in length using Agencourt AMPure XP beads (A63880; Beckman Coulter). Libraries were validated using the High Sensitivity DNA Kit (5067-4626; Agilent) and loaded for pair-end sequencing using Illumina NovaSeq 6000. Trimmomatic tool (Version: 0.39) was used to trim reads with a quality drop below a mean of Q15 in a window of 5 nucleotides and reads with length below 15 nucleotides were filtered out. After the quality control step, the trimmed and filtered reads were aligned to the Zebrafish reference genome GRCz11 using STAR (Version: 2.7.8a) with the parameters “--outFilterMismatchNoverLmax 0.2-outFilterMatchNmin 20 --alignIntronMax 1 -- outFilterMultimapNmax 1” to retain only unique alignments. Reads were deduplicated using Picard (Version: 2.25.0) to remove PCR artefacts. Since the numbers of H3K4me3 peaks may be affected by the sequencing depths, we used the same number of reads (17.5 million pairs) randomly selected from samples of each condition for downstream analysis. The MACS2 peak caller (Version: 2.2.7.1) was employed for each condition with parameters “-q 0.0001 -broad -nomodel - nolambda”. Peaks not located in defined chromosomes were further removed. The filtered peaks were used to do the downstream analysis. Intersection between peaks in CtrlK and DNK conditions was performed with Bedtools toolkit (Version: 2.30.0). Normalized read coverages and subtraction of read coverage were calculated with deeptools toolkit (Version: 2.5.3). ChIPseeker was performed to display the genomic distribution of H3K4me3 peaks based on the matching gene annotation (v. 103) from Ensembl. The H3K4me3 ChIP-seq traces were represented in IGV (Integrative Genomics Viewer) browser. Sequencing data have been deposited in GEO under accession code GSE200937, https://www.ncbi.nlm.nih.gov/geo/query/acc.cgi?acc=GSE200937.

### Quantitative RT-PCR analysis

For FACS-sorted cardiac endothelial cells, RNA from CtrlK sibling and DNK mutant ventricles at 7 dpa was purified using a Magen RNA Nano Kit (R4125; Magen). About 20 ng RNA was used for reverse transcription with MALBAC RNA amplification Kit (KT110700424, YIKON GENOMICS) (Chapman et al., 2015). For RNA extraction from whole hearts, a RNeasy Mini Kit (74106; Qiagen) was used to purify RNA and 500 ng RNA was used for reverse transcription with a Prime Script RT Reagent Kit (RR037A; Takara). Quantitative PCR was performed using a TB Green Premix DimerEraser Kit (RR091A; Takara). The primer sequences are listed in Supplementary Table S1.

### Delivery of chemical Notch inhibitors and siRNAs into adult zebrafish heart

siRNAs were encapsulated in nanoparticles and then injected into the pericardial sac as described previously (Diao et al., 2015; Liu et al., 2013; Xiao et al., 2018; Yang et al., 2011). To evaluate the effect of siRNA-mediated rescue on cardiomyocyte proliferation, 10 μl polyethylene glycol-polylactic acid nanoparticle-encapsulated siRNAs was injected into the pericardial sac daily from 2 to 7 dpa. The Notch inhibitors MK-0752 (HY-10974; MCE) and DAPT (A07D5942; Sigma) were first dissolved in DMSO to make a 20 mM stock solution. Before injection, the stock was diluted to the working concentration (30 μM) and 10 μl of diluted inhibitor was injected daily from 4 to 6 dpa. The injected hearts at 7dpa were then collected for subsequent experiments. siRNA sequences for *notch1a*, *notch1b*, *notch2*, *notch3*, and *kdm7aa* are listed in Supplementary Table S2.

### RNAscope and RNA *in situ* hybridization, immunostaining, and histology

RNAscope (Advanced Cell Diagnostics, Hayward, CA) was applied to 10-μm sections from freshly frozen hearts embedded in Optimal Cutting Temperature (OCT) compound (4583; Sakura). Fresh tissue was fixed in 10% pre-chilled neutral buffered formalin in 1 × PBS at 4°C, followed by dehydration, and then treated with RNAscope® hydrogen peroxide (in RNAscope Universal Pretreatment Kit; 322380; ACD) for 10 min at room temperature. The slides were washed with water and incubated with RNAscope Protease IV (in RNAscope Universal Pretreatment Kit; 322380; ACD) for 30 min at room temperature. Then, they were washed 5 times in 1 × PBS, and the RNAscope® 2.5 HD Duplex Detection Kit (322430; ACD) was applied to visualize hybridization signals. Three injured and sham-operated hearts were used for each RNAscope *in situ* hybridization.

RNA *in situ* hybridization was performed on 10-μm sections from fixed frozen hearts embedded in OCT compound. To generate RNA probes, we amplified *notch1a*, *notch1b*, *notch2*, and *notch3* cDNA from regenerating hearts at 7 dpa, blunt-ligated cDNA into a pEASy-Blunt vector, and generated digoxigenin-labeled RNA probes using T7 RNA polymerases. *In situ* hybridization was performed on cryosections of 4% paraformaldehyde-fixed hearts as previously (Liu, Wang, Li, He, & Liu, 2014).

For immunofluorescence staining, adult zebrafish hearts were fixed in 4% paraformaldehyde at room temperature for 2 h, dehydrated, and embedded in paraffin and sectioned at 5 μm. The sections were dewaxed, rehydrated, and washed in 1 × PBS. The antigens were repaired with the citric acid buffer (CW0128S; CWBIO). After washing, the sections were blocked in 10% FBS in PBST (1% Tween 20 in PBS), and then incubated with diluted primary antibodies (1:150-200 in PBST containing 10% FBS) overnight at 4°C. The primary antibodies used for immunofluorescence were anti-Mef2c (HPA005533; Sigma), anti-GFP (A-11122; Invitrogen), anti-PCNA (P8825; Sigma), anti-myosin heavy-chain monoclonal antibody (14-6503-82; eBioscience), and the Brg1 antibody, which was raised against a glutathione S-transferase-BRG1 fusion protein (human BRG1 amino-acids 1,086-1,307) (Khavari, Peterson, Tamkun, Mendel, & Crabtree, 1993; Wang et al., 1996). After washing, the sections were incubated with secondary antibodies for 2 h at room temperature. The secondary antibodies (1:300 diluted in PBST containing 10% FBS) were Alexa Fluor 488 goat anti-mouse IgG (A21121; Invitrogen), Alexa Fluor 488 goat anti-rabbit IgG (A11034; Invitrogen), Alexa Fluor 555 goat anti-mouse IgG (A21424; Invitrogen), and Alexa Fluor 555 goat anti-rabbit IgG (A21428; Invitrogen).

RNA and RNAscope *in situ* hybridization was examined under a DM5000B microscope (Leica, Germany); immunofluorescence images were captured on a confocal microscope (LSM510; Carl Zeiss, Germany); and fluorescence intensity was quantified using MBF ImageJ.

### Acid fuchsin orange G-stain (AFOG)

AFOG staining was performed on paraffin sections following the manufacturer’s instructions (Han et al., 2014). The sections were incubated in Bouin’s solution (HT10132; Sigma) at 56°C for 2.5 h, and at room temperature for 1 h, washed in tap water, incubated in 1% phosphomolybdic acid (P4869; Sigma) for 5 min, washed with water, and then stained with AFOG solution consisting of 3 g acid fuchsin (F8129; Sigma), 2 g orange G (O3756, Sigma), and 1 g aniline blue (AB0083; BBI) dissolved in 200 ml acidified distilled water (pH 1.1) for 10 min. The sections were rinsed with distilled water, dehydrated, mounted, and staining was photographed under a DM5000B microscope (Leica, Germany).

### Chromatin immunoprecipitation (ChIP) and quantitative ChIP (qChIP)

About 25 zebrafish hearts were pooled for each ChIP experiment. The hearts were dissected from adult zebrafish, and the outflow tract and atrium were removed. Chromatin isolation and ChIP assays were performed using a Pierce Magnetic ChIP Kit (26157; Pierce). Anti-Brg1 (Khavari et al., 1993; Wang et al., 1996) and anti-H3K4me3 (Ab8580, Abcam) antibodies were used for the ChIP assays. The DNA bound by ChIP was used for library construction and quantitative PCR. The primer sequences are listed in Supplementary Table S2.

### Immunoprecipitation (IP)

The full-length coding cDNA of zebrafish *kdm7aa* was isolated from the regenerating heart cDNA library and cloned into the pcDNA3.1 vector. For co-IP, 293T cells (CL-0005, Procell) were transfected with pcDNA3.1-Flag-*kdm7aa*-Myc, pcDNA3.1-Flag-Brg1, and/or pcDNA3.1-Flag-DN-xBrg1 plasmids using Fugen HD Transfection Reagents (2311; Promega), and after 48 h the transfected cells were lysed in NP-40 lysis buffer (P0013F; Beyotime). After brief centrifugation, the supernatants were collected for immunoprecipitation while a protein extraction fraction was set aside for input controls. Equal volumes of supernatants were incubated overnight with 5 μg of either anti-Brg1, anti-Myc, or IgG. Next morning, 25 μl of Pierce Protein A/G Magnetic Beads (88802; Pierce) were added and incubated with the IP mixture for 2 h at room temperature. The beads were then washed for 5 min and repeated 3 times in IP wash buffer (30 mM HEPES, 100 mM NaCl, 1 mM EDTA, 0.5% NP-40, pH 7.5), and were subsequently eluted with 1× loading buffer with heating at 100°C for 10 min. The antibodies for IP were anti-Myc (AT0023; Engibody), anti-Flag (AT0022; Engibody), and anti-Brg1 (J1) (Wang et al., 1996).

### Notch promoter luciferase assays

The promoter sequences of Notch receptors were cloned into the luciferase reporter vector pGL4.26, with the *notch1a* promoter (from 171 bp to +3 bp), *notch1b* promoter (from –41 bp to +58 bp), *notch2* promoter (from –263 bp to –115 bp), and *notch3* promoter (from +394 bp to +504 bp), of which the ATG was considered to be +1 bp. Stable 293T cell lines (CL-0005, Procell) for each of the four notch reporters were generated in the presence of 150 μg/ml hygromycin B. Isolated reporter cells for each of the Notch receptors were co-transfected with pcDNA3.1-*brg1*, pcDNA3.1-*kdm7aa*, pcDNA3.1-DN-*xbrg1*, and pREP4-*Renilla*. Luciferase assays were carried out at 48 h after infection following the manufacturer’s instructions with the Dual-luciferase Reporter Assay System (E1910; Promega). Firefly luciferase activity was normalized by *Renilla* luciferase activity.

### Statistical analysis

All statistics were calculated using Statistical Product and Service Solutions (SPSS) software or GraphPad Prism. The statistical significance of differences between two groups was determined using the independent unpaired *t*-test, with two-tailed P values, and the data are reported as the mean ± s.e.m. Among three or more groups, one-way analysis of variance followed by Bonferroni’s multiple comparison test or Dunnett’s multiple comparison test was used for comparisons.

---

**Figure 1-figure supplement 1.**
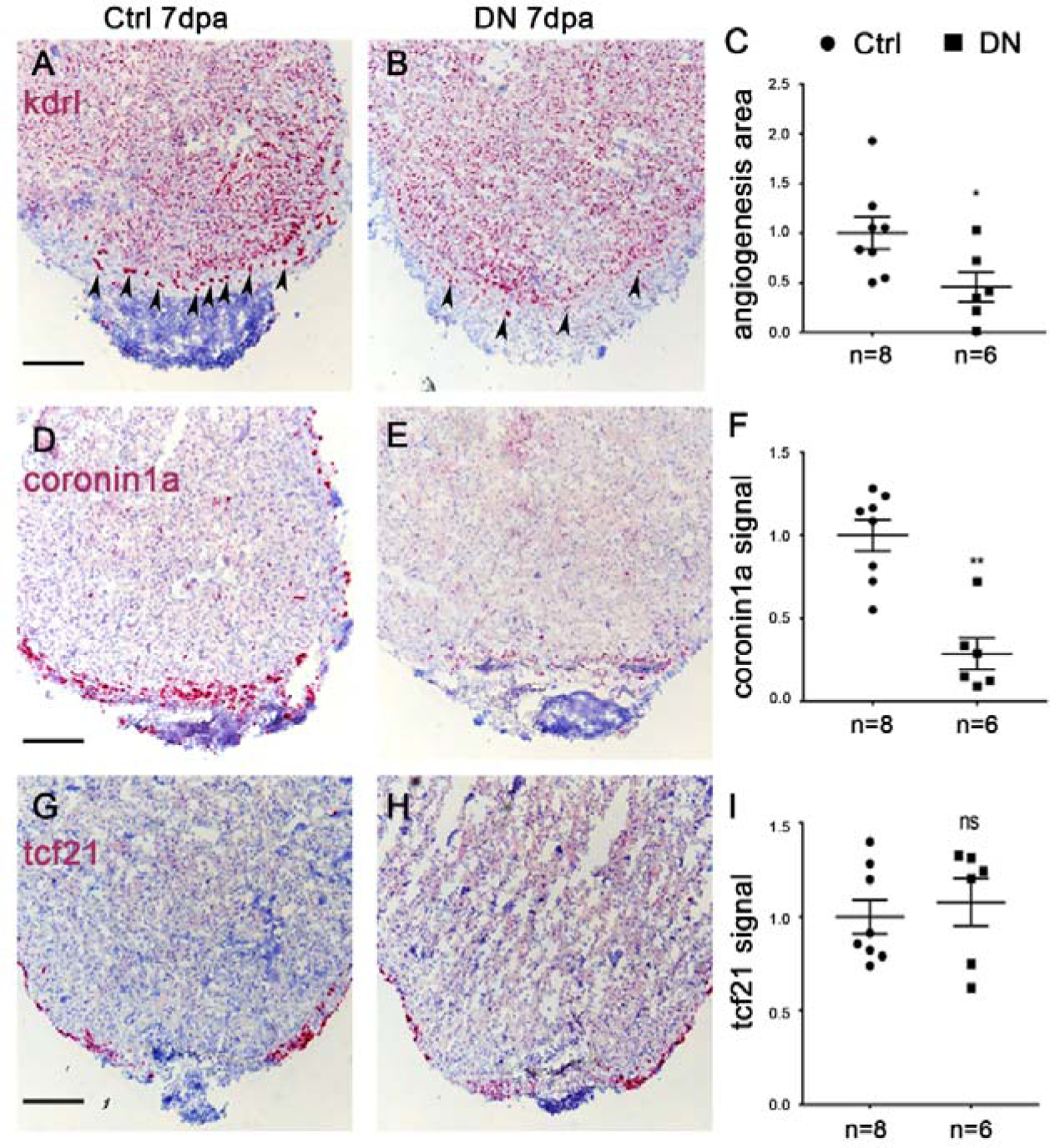
Endothelium-specific inhibition of Brg1 impairs angiogenesis and immune responses but not epicardial activation. **(A, B, D, E, G, H)** RNAscope *in situ* hybridization on representative sections of control sibling hearts [Ctrl: Tg(*ubi*:loxp-DsRed-STOP-loxp-DN-xBrg1)] (A, D, G) and DN-xBrg1 mutant hearts [DN: Tg(*ubi*:loxp-DsRed-STOP-loxp-DN-xBrg1; *kdrl*:CreER) (B, E, H) at 7 dpa, using *kdrl* (endothelial cell marker) (A-B), *coronin1a* (leukocyte marker) (D-E), and *tcf21* (epicardium marker) probes (G-H). Note that endothelium-specific inhibition of Brg1 interferes with *kdrl*-positive endothelial cells (arrowheads) and *coronin1a*-positive leukocyte recruitment while having no effects on *tcf21*-positive epicardium in the presence of 4-HT (scale bars, 100 μm). **(C, F, I)** Statistics of panels A and B (C), D and E (F), and G and H (I). Data are the mean ± s.e.m; *p <0.05, **p <0.01, ns, not significant, unpaired *t*-test. N number shown here (C, F, I) indicate biological repetition. **Figure 1-figure supplement 1-source data 1.** Source data for Figure 1-figure supplement 1A, B. **Figure 1-figure supplement 1-source data 2.** Source data for Figure 1-figure supplement 1D, E. **Figure 1-figure supplement 1-source data 3.** Source data for Figure 1-figure supplement 1G, H. **Figure 1-figure supplement 1-source data 4.** Source data for Figure 1-figure supplement 1C, F, I.

**Figure 2-figure supplement 1.**
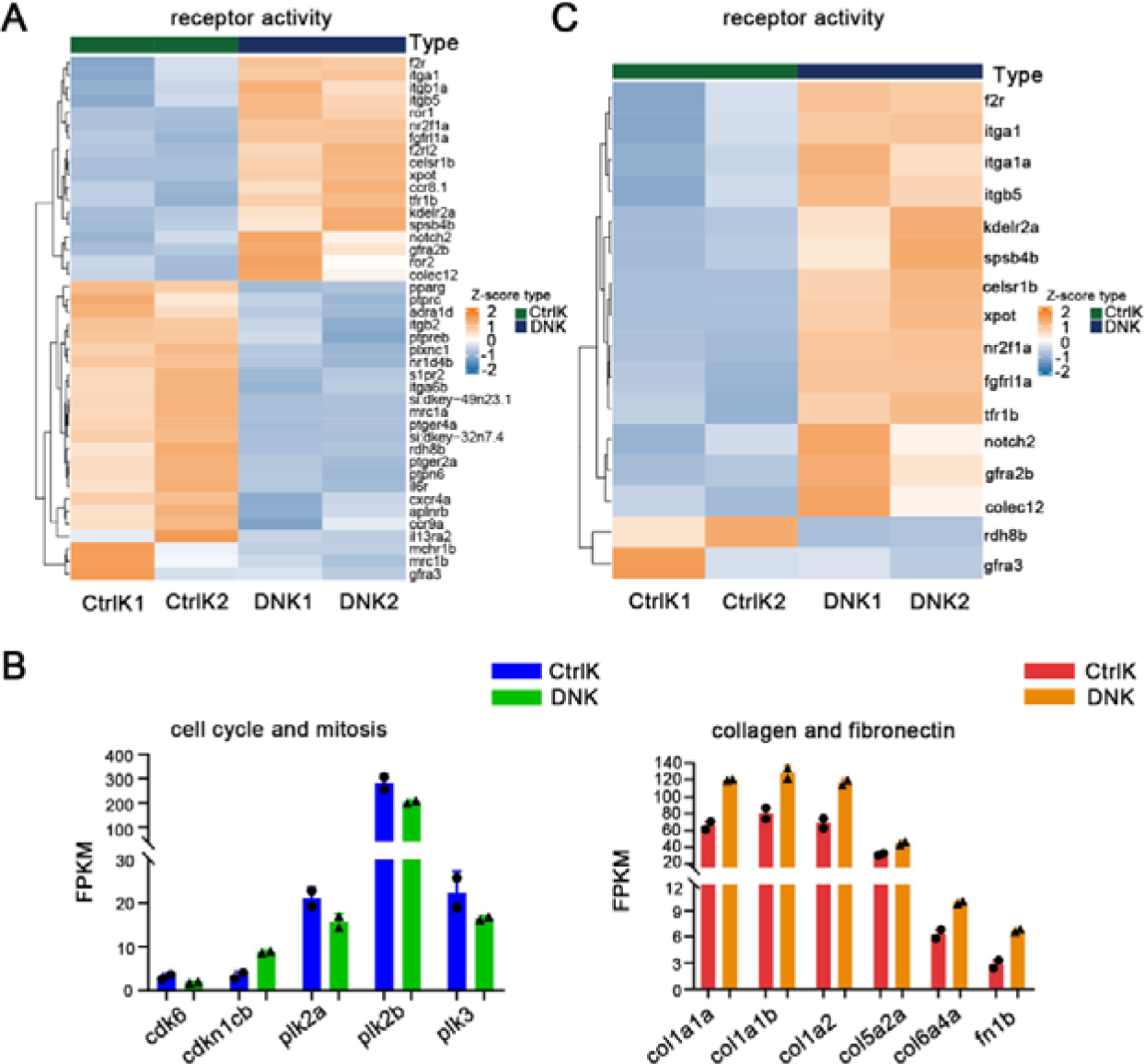
RNA-sequencing analysis shows that endothelial-specific inhibition of Brg1 affects a cluster of receptor genes expression. **(A)** Heat map displaying Z-score normalized gene expressions for receptor activity related genes (GO:0004872) which are differentially expressed (adjusted P-value < 0.05 from DEseq2 result) in FACS-sorted *kdrl*-eGFP positive endothelial cells between Tg(*ubi*:loxp-DsRed-STOP-loxp-DN-xBrg1; *kdrl*:CreER; *kdrl*:eGFP) dominant-negative Brg1 hearts (DNK1 and DNK2) and Tg(*ubi*:loxp-DsRed-STOP-loxp-DN-xBrg1; *kdrl*:eGFP) control hearts (CtrlK1 and CtrlK2) at 7 dpa in the presence of 4-HT. Columns represent individual samples (two biological replicates for each condition). **(B)** Bar graph displaying FPKM values (Fragments Per Kilobase of transcript per Million mapped reads) of representative genes from mitosis and cell cycle, collagen and fibronectin pathways which are down-regulated/up-regulated (adjusted P-value < 0.05 from DEseq2 result) in FACS-sorted *kdrl*-eGFP positive cells from dominant-negative Brg1 hearts (DNK) compared to control hearts (CtrlK). Error bar was indicated by two biological replicates. **(C)** Heat map showing expression of receptor activity related genes that were not only differentially expressed in FACS-sorted *kdrl*-eGFP positive cells between DNK and CtrlK hearts, but were also marked by differentially occupied H3K4me3 peaks in their promoters. **Figure 2-figure supplement 1-source data 1.** Source data for Figure 2-figure supplement 1B.

**Figure 3-figure supplement 1.**
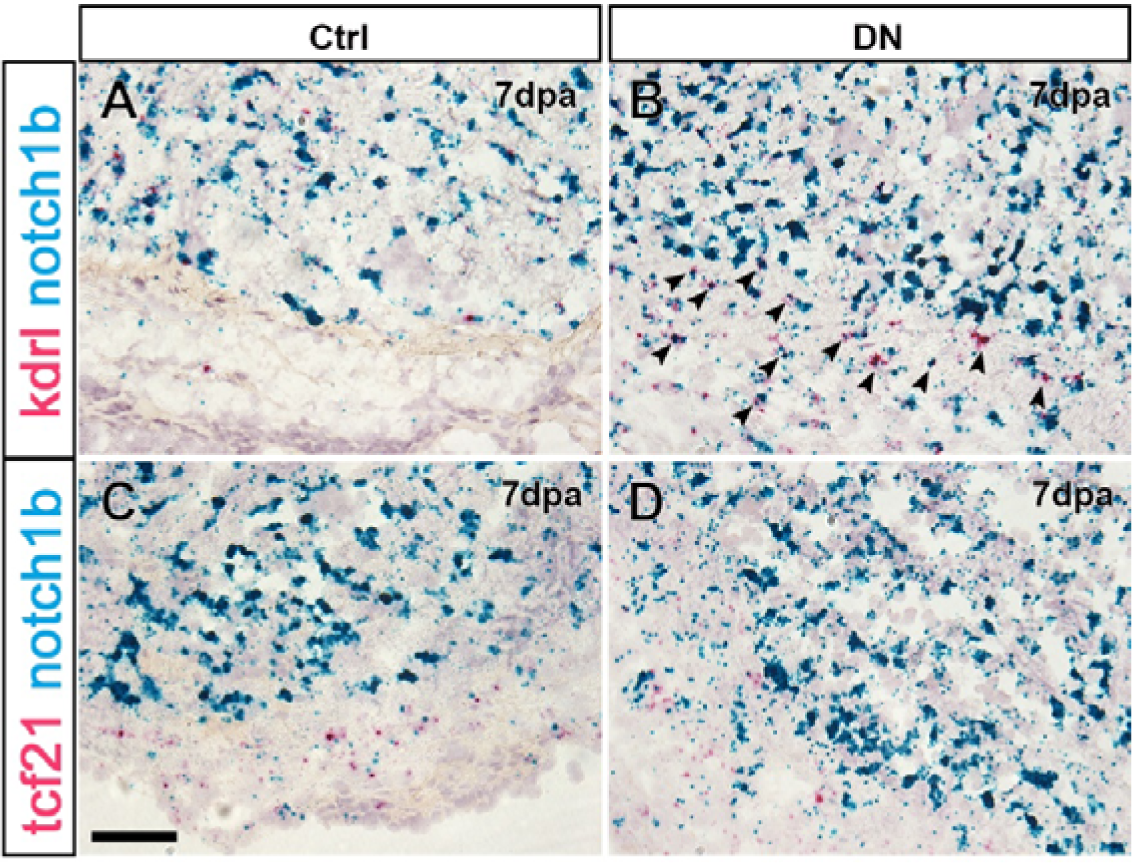
*notch1b* is induced in endothelial cells of hearts expressing DN-xBrg1 after ventricular resection. **(A**–**D)** RNAscope *in situ* hybridization of heart sections from control sibling (Ctrl) (A, C) and dominant-negative (DN) hearts (B, D) at 7 dpa, using either *kdrl* (endothelial cell marker) (A, B) or *tcf21* (epicardium marker) (C, D) probes to co-stain with *notch1b* probes. Note that *notch1b* is particularly induced in *kdrl*-positive endothelial cells, but rarely in *tcf21*-positive epicardium of DN hearts compared with Ctrl hearts in the presence of 4-HT (arrowheads, double-positive signals for both *kdrl* and *notch1b* expression; scale bar, 100 μm).

**Supplementary Table S1.**
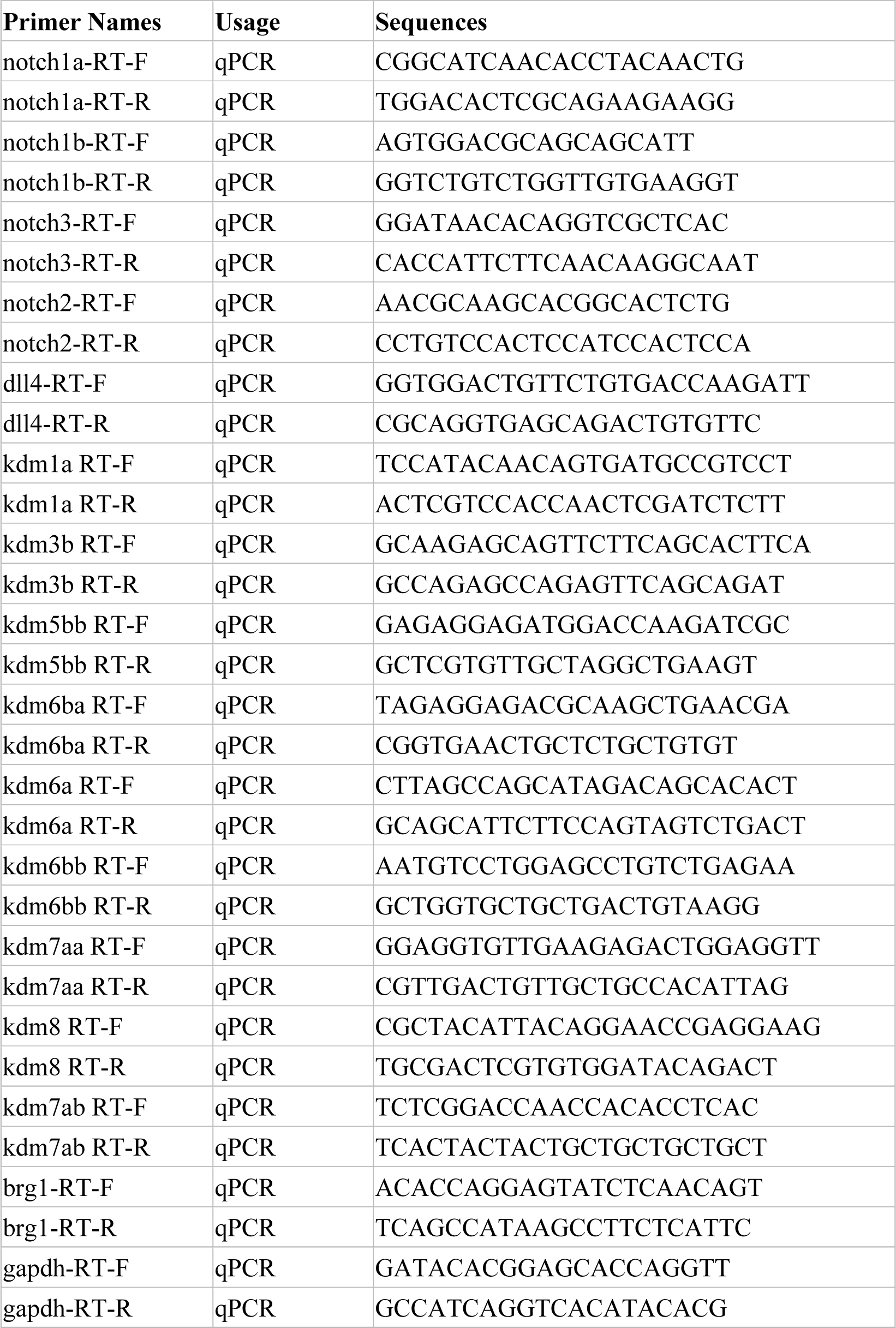

**Supplementary Table S2.**
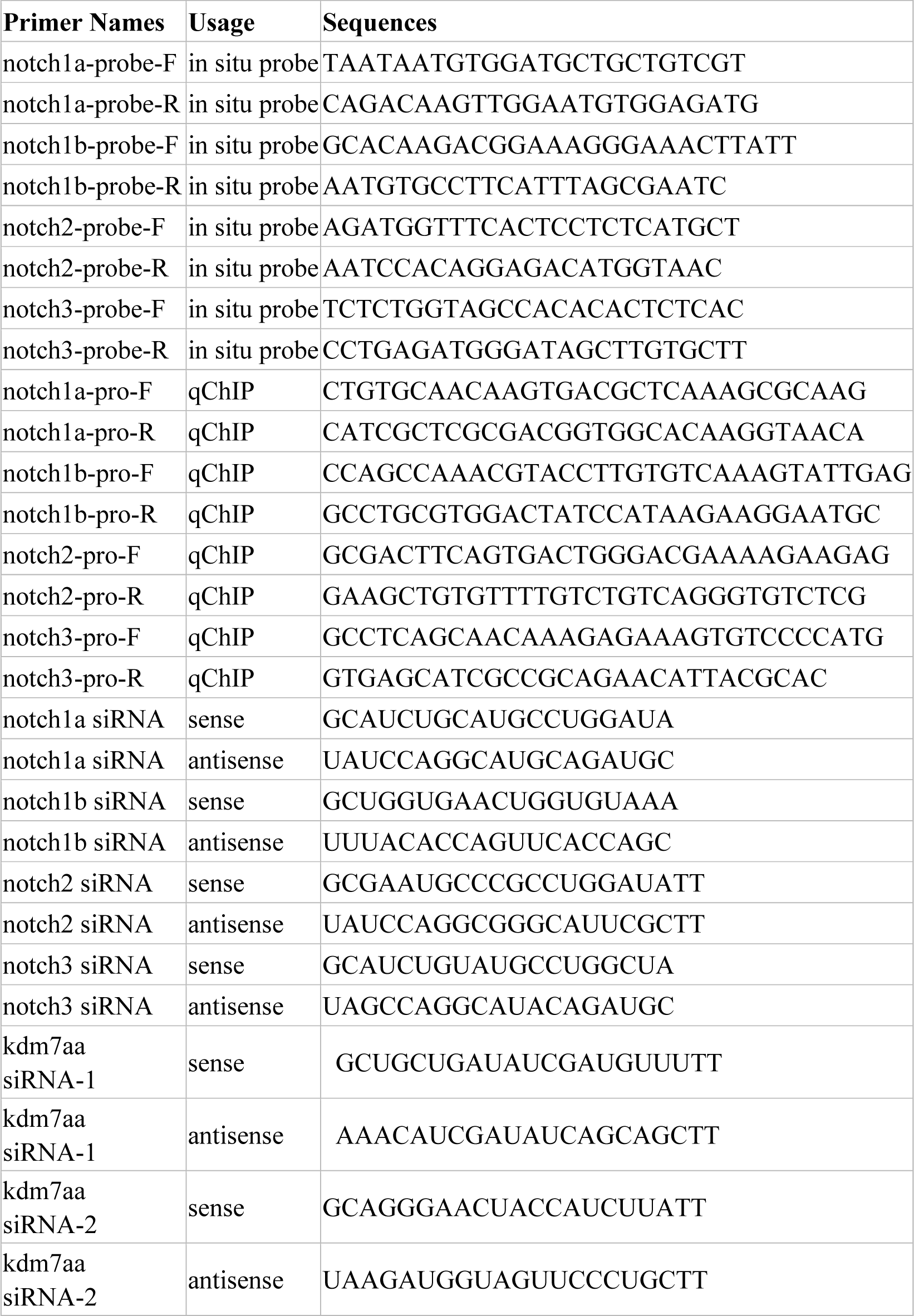

## Notes

### Competing Interest Statement

The authors have declared no competing interest.

## References

1. Beis, D., Bartman, T., Jin, S. W., Scott, I. C., D’Amico, L. A., Ober, E. A., … Jungblut, B. (2005). Genetic and cellular analyses of zebrafish atrioventricular cushion and valve development. Development, 132(18), 4193–4204. Retrieved from https://www.ncbi.nlm.nih.gov/pubmed/16107477. doi:10.1242/dev.01970

2. Bultman, S. J., Gebuhr, T. C., & Magnuson, T. (2005). A Brg1 mutation that uncouples ATPase activity from chromatin remodeling reveals an essential role for SWI/SNF-related complexes in beta-globin expression and erythroid development. Genes Dev, 19(23), 2849–2861. Retrieved from https://www.ncbi.nlm.nih.gov/pubmed/16287714. doi:10.1101/gad.1364105

3. Bultman, S. J., Gebuhr, T. C., Pan, H., Svoboda, P., Schultz, R. M., & Magnuson, T. (2006). Maternal BRG1 regulates zygotic genome activation in the mouse. Genes Dev, 20(13), 1744–1754. Retrieved from https://www.ncbi.nlm.nih.gov/pubmed/16818606. doi:10.1101/gad.1435106

4. Chapman, A. R., He, Z., Lu, S., Yong, J., Tan, L., Tang, F., & Xie, X. S. (2015). Single cell transcriptome amplification with MALBAC. PLoS One, 10(3), e0120889. Retrieved from https://www.ncbi.nlm.nih.gov/pubmed/25822772. doi:10.1371/journal.pone.0120889

5. Chi, T. H., Wan, M., Lee, P. P., Akashi, K., Metzger, D., Chambon, P., … Crabtree, G. R. (2003). Sequential roles of Brg, the ATPase subunit of BAF chromatin remodeling complexes, in thymocyte development. Immunity, 19(2), 169–182. Retrieved from https://www.ncbi.nlm.nih.gov/pubmed/12932351. doi:10.1016/s1074-7613(03)00199-7

6. Diao, J., Wang, H., Chang, N., Zhou, X. H., Zhu, X., Wang, J., & Xiong, J. W. (2015). PEG-PLA nanoparticles facilitate siRNA knockdown in adult zebrafish heart. Dev Biol, 406(2), 196–202. Retrieved from https://www.ncbi.nlm.nih.gov/pubmed/26327645. doi:10.1016/j.ydbio.2015.08.020

7. Duncan, E. M., & Sanchez Alvarado, A. (2019). Regulation of Genomic Output and (Pluri)potency in Regeneration. Annu Rev Genet, 53, 327–346. Retrieved from https://www.ncbi.nlm.nih.gov/pubmed/31505134. doi:10.1146/annurev-genet-112618-043733

8. Eroglu, B., Wang, G., Tu, N., Sun, X., & Mivechi, N. F. (2006). Critical role of Brg1 member of the SWI/SNF chromatin remodeling complex during neurogenesis and neural crest induction in zebrafish. Dev Dyn, 235(10), 2722–2735. Retrieved from https://www.ncbi.nlm.nih.gov/pubmed/16894598. doi:10.1002/dvdy.20911

9. Gao, J., Fan, L., Zhao, L., & Su, Y. (2021). The interaction of Notch and Wnt signaling pathways in vertebrate regeneration. Cell Regen, 10(1), 11. Retrieved from https://www.ncbi.nlm.nih.gov/pubmed/33791915. doi:10.1186/s13619-020-00072-2

10. Gemberling, M., Bailey, T. J., Hyde, D. R., & Poss, K. D. (2013). The zebrafish as a model for complex tissue regeneration. Trends Genet, 29(11), 611–620. Retrieved from https://www.ncbi.nlm.nih.gov/pubmed/23927865. doi:10.1016/j.tig.2013.07.003

11. Goldman, J. A., Kuzu, G., Lee, N., Karasik, J., Gemberling, M., Foglia, M. J., … Poss, K. D. (2017). Resolving Heart Regeneration by Replacement Histone Profiling. Dev Cell, 40(4), 392–404 e395. Retrieved from https://www.ncbi.nlm.nih.gov/pubmed/28245924. doi:10.1016/j.devcel.2017.01.013

12. Griffin, C. T., Brennan, J., & Magnuson, T. (2008). The chromatin-remodeling enzyme BRG1 plays an essential role in primitive erythropoiesis and vascular development. Development, 135(3), 493–500. Retrieved from https://www.ncbi.nlm.nih.gov/pubmed/18094026. doi:10.1242/dev.010090

13. Gu, Z., Eils, R., & Schlesner, M. (2016). Complex heatmaps reveal patterns and correlations in multidimensional genomic data. Bioinformatics, 32(18), 2847–2849. Retrieved from https://www.ncbi.nlm.nih.gov/pubmed/27207943. doi:10.1093/bioinformatics/btw313

14. Han, P., Zhou, X. H., Chang, N., Xiao, C. L., Yan, S., Ren, H., … Xiong, J. W. (2014). Hydrogen peroxide primes heart regeneration with a derepression mechanism. Cell Res, 24(9), 1091–1107. Retrieved from https://www.ncbi.nlm.nih.gov/pubmed/25124925. doi:10.1038/cr.2014.108

15. Hang, C. T., Yang, J., Han, P., Cheng, H. L., Shang, C., Ashley, E., … Chang, C. P. (2010). Chromatin regulation by Brg1 underlies heart muscle development and disease. Nature, 466(7302), 62–67. Retrieved from https://www.ncbi.nlm.nih.gov/pubmed/20596014. doi:10.1038/nature09130

16. Harikumar, A., & Meshorer, E. (2015). Chromatin remodeling and bivalent histone modifications in embryonic stem cells. EMBO Rep, 16(12), 1609–1619. Retrieved from https://www.ncbi.nlm.nih.gov/pubmed/26553936. doi:10.15252/embr.201541011

17. Hesse, M., Welz, A., & Fleischmann, B. K. (2018). Heart regeneration and the cardiomyocyte cell cycle. Pflugers Arch, 470(2), 241–248. Retrieved from https://www.ncbi.nlm.nih.gov/pubmed/28849267. doi:10.1007/s00424-017-2061-4

18. Ho, L., & Crabtree, G. R. (2010). Chromatin remodelling during development. Nature, 463(7280), 474–484. Retrieved from https://www.ncbi.nlm.nih.gov/pubmed/20110991. doi:10.1038/nature08911

19. Jopling, C., Sleep, E., Raya, M., Marti, M., Raya, A., & Izpisua Belmonte, J. C. (2010). Zebrafish heart regeneration occurs by cardiomyocyte dedifferentiation and proliferation. Nature, 464(7288), 606–609. Retrieved from https://www.ncbi.nlm.nih.gov/pubmed/20336145. doi:10.1038/nature08899

20. Kawakami, K., Takeda, H., Kawakami, N., Kobayashi, M., Matsuda, N., & Mishina, M. (2004). A transposon-mediated gene trap approach identifies developmentally regulated genes in zebrafish. Dev Cell, 7(1), 133–144. Retrieved from https://www.ncbi.nlm.nih.gov/pubmed/15239961. doi:10.1016/j.devcel.2004.06.005

21. Khavari, P. A., Peterson, C. L., Tamkun, J. W., Mendel, D. B., & Crabtree, G. R. (1993). BRG1 contains a conserved domain of the SWI2/SNF2 family necessary for normal mitotic growth and transcription. Nature, 366(6451), 170–174. Retrieved from https://www.ncbi.nlm.nih.gov/pubmed/8232556. doi:10.1038/366170a0

22. Kikuchi, K., Holdway, J. E., Major, R. J., Blum, N., Dahn, R. D., Begemann, G., & Poss, K. D. (2011). Retinoic acid production by endocardium and epicardium is an injury response essential for zebrafish heart regeneration. Dev Cell, 20(3), 397–404. Retrieved from https://www.ncbi.nlm.nih.gov/pubmed/21397850. doi:10.1016/j.devcel.2011.01.010

23. Kikuchi, K., Holdway, J. E., Werdich, A. A., Anderson, R. M., Fang, Y., Egnaczyk, G. F., … Poss, K. D. (2010). Primary contribution to zebrafish heart regeneration by gata4(+) cardiomyocytes. Nature, 464(7288), 601–605. Retrieved from https://www.ncbi.nlm.nih.gov/pubmed/20336144. doi:10.1038/nature08804

24. Kim, D., Paggi, J. M., Park, C., Bennett, C., & Salzberg, S. L. (2019). Graph-based genome alignment and genotyping with HISAT2 and HISAT-genotype. Nat Biotechnol, 37(8), 907–915. Retrieved from https://www.ncbi.nlm.nih.gov/pubmed/31375807. doi:10.1038/s41587-019-0201-4

25. Li, G., & Reinberg, D. (2011). Chromatin higher-order structures and gene regulation. Curr Opin Genet Dev, 21(2), 175–186. Retrieved from https://www.ncbi.nlm.nih.gov/pubmed/21342762. doi:10.1016/j.gde.2011.01.022

26. Li, N., Kong, M., Zeng, S., Hao, C., Li, M., Li, L., … Xu, Y. (2019). Brahma related gene 1 (Brg1) contributes to liver regeneration by epigenetically activating the Wnt/beta-catenin pathway in mice. FASEB J, 33(1), 327–338. Retrieved from https://www.ncbi.nlm.nih.gov/pubmed/30001167. doi:10.1096/fj.201800197R

27. Liao, Y., Smyth, G. K., & Shi, W. (2014). featureCounts: an efficient general purpose program for assigning sequence reads to genomic features. Bioinformatics, 30(7), 923–930. Retrieved from https://www.ncbi.nlm.nih.gov/pubmed/24227677. doi:10.1093/bioinformatics/btt656

28. Liu, C. C., Sun, C., Zheng, X., Zhao, M. Q., Kong, F., Xu, F. L., … Xia, M. (2019). Regulation of KDM2B and Brg1 on Inflammatory Response of Nasal Mucosa in CRSwNP. Inflammation, 42(4), 1389–1400. Retrieved from https://www.ncbi.nlm.nih.gov/pubmed/31041569. doi:10.1007/s10753-019-01000-6

29. Liu, J., Gu, C., Cabigas, E. B., Pendergrass, K. D., Brown, M. E., Luo, Y., & Davis, M. E. (2013). Functionalized dendrimer-based delivery of angiotensin type 1 receptor siRNA for preserving cardiac function following infarction. Biomaterials, 34(14), 3729–3736. Retrieved from https://www.ncbi.nlm.nih.gov/pubmed/23433774. doi:10.1016/j.biomaterials.2013.02.008

30. Liu, K. L., Wang, X. M., Li, Z. L., He, R. Q., & Liu, Y. (2014). In situ hybridization and immunostaining of Xenopus brain. Methods Mol Biol, 1082, 129–141. Retrieved from https://www.ncbi.nlm.nih.gov/pubmed/24048931. doi:10.1007/978-1-62703-655-9_9

31. Love, M. I., Huber, W., & Anders, S. (2014). Moderated estimation of fold change and dispersion for RNA-seq data with DESeq2. Genome Biol, 15(12), 550. Retrieved from https://www.ncbi.nlm.nih.gov/pubmed/25516281. doi:10.1186/s13059-014-0550-8

32. Martinez-Redondo, P., & Izpisua Belmonte, J. C. (2020). Tailored chromatin modulation to promote tissue regeneration. Semin Cell Dev Biol, 97, 3–15. Retrieved from https://www.ncbi.nlm.nih.gov/pubmed/31028854. doi:10.1016/j.semcdb.2019.04.015

33. Menon, D. U., Shibata, Y., Mu, W., & Magnuson, T. (2019). Mammalian SWI/SNF collaborates with a polycomb-associated protein to regulate male germline transcription in the mouse. Development, 146(19). Retrieved from https://www.ncbi.nlm.nih.gov/pubmed/31043422. doi:10.1242/dev.174094

34. Mosimann, C., Kaufman, C. K., Li, P., Pugach, E. K., Tamplin, O. J., & Zon, L. I. (2011). Ubiquitous transgene expression and Cre-based recombination driven by the ubiquitin promoter in zebrafish. Development, 138(1), 169–177. Retrieved from https://www.ncbi.nlm.nih.gov/pubmed/21138979. doi:10.1242/dev.059345

35. Munch, J., Grivas, D., Gonzalez-Rajal, A., Torregrosa-Carrion, R., & de la Pompa, J. L. (2017). Notch signalling restricts inflammation and serpine1 expression in the dynamic endocardium of the regenerating zebrafish heart. Development, 144(8), 1425–1440. Retrieved from https://www.ncbi.nlm.nih.gov/pubmed/28242613. doi:10.1242/dev.143362

36. Myers, T. R., Amendola, P. G., Lussi, Y. C., & Salcini, A. E. (2018). JMJD-1.2 controls multiple histone post-translational modifications in germ cells and protects the genome from replication stress. Sci Rep, 8(1), 3765. Retrieved from https://www.ncbi.nlm.nih.gov/pubmed/29491442. doi:10.1038/s41598-018-21914-9

37. Oyama, K., El-Nachef, D., Zhang, Y., Sdek, P., & MacLellan, W. R. (2014). Epigenetic regulation of cardiac myocyte differentiation. Front Genet, 5, 375. Retrieved from https://www.ncbi.nlm.nih.gov/pubmed/25408700. doi:10.3389/fgene.2014.00375

38. Patra, C., Kontarakis, Z., Kaur, H., Rayrikar, A., Mukherjee, D., & Stainier, D. Y. R. (2017). The zebrafish ventricle: A hub of cardiac endothelial cells for in vitro cell behavior studies. Sci Rep, 7(1), 2687. Retrieved from https://www.ncbi.nlm.nih.gov/pubmed/28578380. doi:10.1038/s41598-017-02461-1

39. Pertea, M., Pertea, G. M., Antonescu, C. M., Chang, T. C., Mendell, J. T., & Salzberg, S. L. (2015). StringTie enables improved reconstruction of a transcriptome from RNA-seq reads. Nat Biotechnol, 33(3), 290–295. Retrieved from https://www.ncbi.nlm.nih.gov/pubmed/25690850. doi:10.1038/nbt.3122

40. Porrello, E. R., Mahmoud, A. I., Simpson, E., Hill, J. A., Richardson, J. A., Olson, E. N., & Sadek, H. A. (2011). Transient regenerative potential of the neonatal mouse heart. Science, 331(6020), 1078–1080. Retrieved from https://www.ncbi.nlm.nih.gov/pubmed/21350179. doi:10.1126/science.1200708

41. Poss, K. D., Wilson, L. G., & Keating, M. T. (2002). Heart regeneration in zebrafish. Science, 298(5601), 2188–2190. Retrieved from https://www.ncbi.nlm.nih.gov/pubmed/12481136. doi:10.1126/science.1077857

42. Pronobis, M. I., & Poss, K. D. (2020). Signals for cardiomyocyte proliferation during zebrafish heart regeneration. Curr Opin Physiol, 14, 78–85. Retrieved from https://www.ncbi.nlm.nih.gov/pubmed/32368708. doi:10.1016/j.cophys.2020.02.002

43. Quinlan, A. R., & Hall, I. M. (2010). BEDTools: a flexible suite of utilities for comparing genomic features. Bioinformatics, 26(6), 841–842. Retrieved from https://www.ncbi.nlm.nih.gov/pubmed/20110278. doi:10.1093/bioinformatics/btq033

44. Ramirez, F., Ryan, D. P., Gruning, B., Bhardwaj, V., Kilpert, F., Richter, A. S., … Manke, T. (2016). deepTools2: a next generation web server for deep-sequencing data analysis. Nucleic Acids Res, 44(W1), W160–165. Retrieved from https://www.ncbi.nlm.nih.gov/pubmed/27079975. doi:10.1093/nar/gkw257

45. Raya, A., Koth, C. M., Buscher, D., Kawakami, Y., Itoh, T., Raya, R. M., … Izpisua-Belmonte, J. C. (2003). Activation of Notch signaling pathway precedes heart regeneration in zebrafish. Proc Natl Acad Sci U S A, 100 *Suppl 1*, 11889–11895. Retrieved from https://www.ncbi.nlm.nih.gov/pubmed/12909711. doi:10.1073/pnas.1834204100

46. Robinson, J. T., Thorvaldsdottir, H., Winckler, W., Guttman, M., Lander, E. S., Getz, G., & Mesirov, J. P. (2011). Integrative genomics viewer. Nat Biotechnol, 29(1), 24–26. Retrieved from https://www.ncbi.nlm.nih.gov/pubmed/21221095. doi:10.1038/nbt.1754

47. Roman, B. L., Pham, V. N., Lawson, N. D., Kulik, M., Childs, S., Lekven, A. C., … Weinstein, B. M. (2002). Disruption of acvrl1 increases endothelial cell number in zebrafish cranial vessels. Development, 129(12), 3009–3019. Retrieved from https://www.ncbi.nlm.nih.gov/pubmed/12050147.

48. Sadek, H., & Olson, E. N. (2020). Toward the Goal of Human Heart Regeneration. Cell Stem Cell, 26(1), 7–16. Retrieved from https://www.ncbi.nlm.nih.gov/pubmed/31901252. doi:10.1016/j.stem.2019.12.004

49. Schneider, C. A., Rasband, W. S., & Eliceiri, K. W. (2012). NIH Image to ImageJ: 25 years of image analysis. Nat Methods, 9(7), 671–675. Retrieved from https://www.ncbi.nlm.nih.gov/pubmed/22930834. doi:10.1038/nmeth.2089

50. Seo, S., Richardson, G. A., & Kroll, K. L. (2005). The SWI/SNF chromatin remodeling protein Brg1 is required for vertebrate neurogenesis and mediates transactivation of Ngn and NeuroD. Development, 132(1), 105–115. Retrieved from https://www.ncbi.nlm.nih.gov/pubmed/15576411. doi:10.1242/dev.01548

51. Stankunas, K., Hang, C. T., Tsun, Z. Y., Chen, H., Lee, N. V., Wu, J. I., … Chang, C. P. (2008). Endocardial Brg1 represses ADAMTS1 to maintain the microenvironment for myocardial morphogenesis. Dev Cell, 14(2), 298–311. Retrieved from https://www.ncbi.nlm.nih.gov/pubmed/18267097. doi:10.1016/j.devcel.2007.11.018

52. Stewart, S., Tsun, Z. Y., & Izpisua Belmonte, J. C. (2009). A histone demethylase is necessary for regeneration in zebrafish. Proc Natl Acad Sci U S A, 106(47), 19889–19894. Retrieved from https://www.ncbi.nlm.nih.gov/pubmed/19897725. doi:10.1073/pnas.0904132106

53. Tsukada, Y., Ishitani, T., & Nakayama, K. I. (2010). KDM7 is a dual demethylase for histone H3 Lys 9 and Lys 27 and functions in brain development. Genes Dev, 24(5), 432–437. Retrieved from https://www.ncbi.nlm.nih.gov/pubmed/20194436. doi:10.1101/gad.1864410

54. Tzahor, E., & Poss, K. D. (2017). Cardiac regeneration strategies: Staying young at heart. Science, 356(6342), 1035–1039. Retrieved from https://www.ncbi.nlm.nih.gov/pubmed/28596337. doi:10.1126/science.aam5894

55. Vastenhouw, N. L., Zhang, Y., Woods, I. G., Imam, F., Regev, A., Liu, X. S., … Schier, A. F. (2010). Chromatin signature of embryonic pluripotency is established during genome activation. Nature, 464(7290), 922–926. Retrieved from https://www.ncbi.nlm.nih.gov/pubmed/20336069. doi:10.1038/nature08866

56. Wang, W., Cote, J., Xue, Y., Zhou, S., Khavari, P. A., Biggar, S. R., … Crabtree, G. R. (1996). Purification and biochemical heterogeneity of the mammalian SWI-SNF complex. EMBO J, 15(19), 5370–5382. Retrieved from https://www.ncbi.nlm.nih.gov/pubmed/8895581.

57. Wang, W., Hu, C. K., Zeng, A., Alegre, D., Hu, D., Gotting, K., … Sanchez Alvarado, A. (2020). Changes in regeneration-responsive enhancers shape regenerative capacities in vertebrates. Science, 369(6508). Retrieved from https://www.ncbi.nlm.nih.gov/pubmed/32883834. doi:10.1126/science.aaz3090

58. Xiao, C., Gao, L., Hou, Y., Xu, C., Chang, N., Wang, F., … Xiong, J. W. (2016). Chromatin-remodelling factor Brg1 regulates myocardial proliferation and regeneration in zebrafish. Nat Commun, 7, 13787. Retrieved from https://www.ncbi.nlm.nih.gov/pubmed/27929112. doi:10.1038/ncomms13787

59. Xiao, C., Wang, F., Hou, J., Zhu, X., Luo, Y., & Xiong, J. W. (2018). Nanoparticle-mediated siRNA Gene-silencing in Adult Zebrafish Heart. J Vis Exp(137). Retrieved from https://www.ncbi.nlm.nih.gov/pubmed/30102293. doi:10.3791/58054

60. Yang, X. Z., Dou, S., Sun, T. M., Mao, C. Q., Wang, H. X., & Wang, J. (2011). Systemic delivery of siRNA with cationic lipid assisted PEG-PLA nanoparticles for cancer therapy. J Control Release, 156(2), 203–211. Retrieved from https://www.ncbi.nlm.nih.gov/pubmed/21839126. doi:10.1016/j.jconrel.2011.07.035

61. Yu, G., Wang, L. G., & He, Q. Y. (2015). ChIPseeker: an R/Bioconductor package for ChIP peak annotation, comparison and visualization. Bioinformatics, 31(14), 2382–2383. Retrieved from https://www.ncbi.nlm.nih.gov/pubmed/25765347. doi:10.1093/bioinformatics/btv145

62. Zhan, Y., Huang, Y., Chen, J., Cao, Z., He, J., Zhang, J., … Li, L. (2018). The caudal dorsal artery generates hematopoietic stem and progenitor cells via the endothelial-to-hematopoietic transition in zebrafish. J Genet Genomics. Retrieved from https://www.ncbi.nlm.nih.gov/pubmed/29929848. doi:10.1016/j.jgg.2018.02.010

63. Zhang, Y., Yuan, Y., Li, Z., Chen, H., Fang, M., Xiao, P., & Xu, Y. (2019). An interaction between BRG1 and histone modifying enzymes mediates lipopolysaccharide-induced proinflammatory cytokines in vascular endothelial cells. J Cell Biochem, 120(8), 13216–13225. Retrieved from https://www.ncbi.nlm.nih.gov/pubmed/30891798. doi:10.1002/jcb.28595

64. Zhao, L., Ben-Yair, R., Burns, C. E., & Burns, C. G. (2019). Endocardial Notch Signaling Promotes Cardiomyocyte Proliferation in the Regenerating Zebrafish Heart through Wnt Pathway Antagonism. Cell Rep, 26(3), 546–554 e545. Retrieved from https://www.ncbi.nlm.nih.gov/pubmed/30650349. doi:10.1016/j.celrep.2018.12.048

65. Zhao, L., Borikova, A. L., Ben-Yair, R., Guner-Ataman, B., MacRae, C. A., Lee, R. T., … Burns, C. E. (2014). Notch signaling regulates cardiomyocyte proliferation during zebrafish heart regeneration. Proc Natl Acad Sci U S A, 111(4), 1403–1408. Retrieved from https://www.ncbi.nlm.nih.gov/pubmed/24474765. doi:10.1073/pnas.1311705111

66. Zheng, L., Du, J., Wang, Z., Zhou, Q., Zhu, X., & Xiong, J. W. (2021). Molecular regulation of myocardial proliferation and regeneration. Cell Regen, 10(1), 13. Retrieved from https://www.ncbi.nlm.nih.gov/pubmed/33821373. doi:10.1186/s13619-021-00075-7

67. Zhu, W., Xu, X., Wang, X., & Liu, J. (2019). Reprogramming histone modification patterns to coordinate gene expression in early zebrafish embryos. BMC Genomics, 20(1), 248. Retrieved from https://www.ncbi.nlm.nih.gov/pubmed/30922236. doi:10.1186/s12864-019-5611-7

68. Zhu, X., Xiao, C., & Xiong, J. W. (2018). Epigenetic Regulation of Organ Regeneration in Zebrafish. J Cardiovasc Dev Dis, 5(4). Retrieved from https://www.ncbi.nlm.nih.gov/pubmed/30558240. doi:10.3390/jcdd5040057

